# The compartmentalized activity of a legume subtilase SBT12a allows symbiosome stabilization

**DOI:** 10.64898/2026.04.09.717438

**Authors:** Guofeng Zhang, Fabian Stockert, Melissa Mantz, Marta Rodriguez-Franco, Fabian van Beveren, Casandra Hernández-Reyes, Beatrice Lace, Wei Yang, Chris van der Does, Hector Mancilla, Aida Maric, Nils Nebel, Sjon Hartman, Sonja-Verena Albers, Claudine Kraft, Chao Su, Pierre-Marc Delaux, Pitter F. Huesgen, Thomas Ott

## Abstract

The stabilization of rhizobia in specialized organelles, called symbiosomes, is an evolutionary hallmark for maintaining high rates of nitrogen fixation in legumes. This is achieved by releasing thousands of bacteria from nodular infection threads into a single plant cell, which poses a great challenge for the host to keep control over these differentiated bacteria. Considering the importance of proteolytic degradation of antimicrobial proteins or generation of symbiosis-promoting signalling factors, proteolytic activity may represent a key regulatory system in this process. Here, we identified the *Medicago truncatula* subtilisin-like serine protease (subtilase, SBT) 12a acting as a novel regulator of symbiosome functionality and maintenance. Loss-of-function mutations in *SBT12a* led to severe symbiotic defects with nodules of *sbt12a* being characterized by high level induction of defense/senescence-related genes. Using untargeted proteomic High-efficiency Undecanal-based N-Termini EnRichment (HUNTER), we identified candidate substrates of SBT12a and individually confirmed intrinsic cleavage sites. Our analysis revealed that SBT12a activity within symbiosomes is associated with the proteolytic regulation of DNF2, a symbiotic protein that is specifically required to prevent defense response and premature nodule senescence during nitrogen fixation. Together, our findings position SBT12a as a central host factor controlling symbiosome performance.

## Main

The ability of legume plants to form symbiotic associations with nitrogen-fixing rhizobia (root nodule symbiosis; RNS) allows them to overcome nitrogen deficiency in the soil. While this ability has been lost several times independently during evolution (Griesmann et al., 2018), legumes, which maintained this unique mutualism, stabilized their intracellular symbionts in newly formed root organs, called nodules, and further compartmentalized them in specialized cell organelles, known as the symbiosomes. Given their tremendous importance and despite their complex nature, nodule-hosted nitrogen-fixing symbiosomes are considered key targets for engineering RNS in non-legume crops to sustainably reduce the amounts of chemical nitrogen fertilizers in agriculture (Huisman & Geurts, 2020, Jhu & Oldroyd, 2023).

Over the past decades, our understanding of the initial molecular dialogue between soil-borne rhizobia and their host plants has advanced considerably (Ghantasala & Roy Choudhury, 2026, Ruan et al., 2026). The corresponding receptors are able, when being stably associated, to trigger nodule organogenesis in the absence of the symbiont (Rübsam et al., 2023). However, these and other spontaneous nodules have so far never been colonized by rhizobia. In many legumes such as *Medicago truncatula* (hereafter, Medicago), successful infection of the host root and nodules is achieved by the formation of infection threads (ITs) (de Carvalho-Niebel et al., 2024, Gao et al., 2024). To support polar growth of ITs which enables transcellular progression of rhizobia across several cell layers, host cells undergo extensive cellular reprogramming that involves local cell wall modifications, membrane repolarization, cytoskeleton rearrangement and cell cycle regulation (Liang et al., 2018, Gao et al., 2022, Lace et al., 2023, Su et al., 2023, Zhang & Ott, 2024, Batzenschlager et al., 2025, Zhao et al., 2025, Qiao et al., 2026). The ultimate step to stabilize the functional symbiosis is the release of the bacteria from nodular ITs (nITs) into infection-competent nodule cells. This step occurs at so-called infection droplets (IDs), bulged and cell wall-free sites of nITs from where thousands of bacteria are pinched off into a single host cell while remaining encapsulated by a plant-derived membrane forming the nitrogen-fixing symbiosome (Adema & Kohlen, 2024, Zhang et al., 2024). At their peak of symbiotic performance, these infected cells are densely packed with bacteria which, in the case of Medicago belonging to the inverted repeat-lacking clade (IRLC), terminally differentiate into bacteroids. A mature symbiosome thus comprises three distinct compartments: the symbiosome membrane, the bacteroid, and the intervening space termed peribacteroid space. Bacteroid differentiation is hallmarked by genome duplications, membrane permeabilization as well as cell elongation, and substantially controlled by a large plant gene family (over 700 members) encoding NODULE-SPECIFIC CYSTEINE-RICH (NCR) peptides (Alunni & Gourion, 2016, Guerra-Garcia & Sankari, 2025). Although exhibiting antimicrobial activity in vitro, these NCR peptides, which are targeted to symbiosomes via the DEFECTIVE IN NITROGEN FIXATION (DNF)1-mediated nodule-specific protein secretory pathway, were shown to be crucial for bacteroid survival and differentiation *in planta* (Kim et al., 2015, Wang et al., 2010, Van de Velde et al., 2010). For instance, *NCR169* was identified as a single causal gene in the *dnf7* mutant, which displays impaired bacteroid persistence, premature nodule senescence and reduced nitrogen fixation ability (Horváth et al., 2015). Moreover, accumulating evidence suggests that, in addition to NCR peptides, several other plant factors are also indispensable for bacteroid differentiation, including the phosphatidylinositol-specific phospholipase C-like (PI-PLC) X domain protein DNF2 (Bourcy et al., 2013), the SYMBIOTIC CYSTEINE-RICH RECEPTOR-LIKE KINASE (SymCRK) (Berrabah et al., 2014), NODULES WITH ACTIVATED DEFENSE 1 (NAD1) (Wang et al., 2016), REGULATOR OF SYMBIOSOME DIFFERENTIATION (RSD) (Sinharoy et al., 2013), and NODULE-SPECIFIC POLYCYSTIN-1, LIPOEXYGENASE, α-TOXIN (PLAT) DOMAIN (NPD) proteins (Pislariu et al., 2019, Trujillo et al., 2019). Loss-of-function mutations in any of these genes result in severe defects in symbiosome development, disrupted nodule zonation, and the failure to maintain functional nitrogen-fixing nodules. Furthermore, the tight association between the endoplasmic reticulum (ER), symbiosomes and the plasma membrane also poses challenges to fine-tune targeted secretion and distribution of proteins to the symbiosome membrane/space and the plasma membrane/apoplast. This was recently exemplified for the peribacteroid space. Although being cell wall-free, it requires constant clearing of antimicrobial pectin fragments by lytic enzymes that are continuously secreted into the peribacteroid space (Gao et al., 2025).

These processes further raise the demand for proteases that inactivate antimicrobial proteins or generate symbiosis-promoting signalling proteins/peptides in the peribacteroid space. One of the largest families of proteolytic enzymes potentially responsible for such mechanisms are subtilisin-like serine proteases (subtilases, SBTs), which are characterized by a specific arrangement of Asp, His and Ser residues in the catalytic triad (Schaller et al., 2018). Plant SBTs are typically synthesized as preproenzymes comprising an N-terminal signal peptide for secretion, an inhibitory prodomain that is self-cleaved during maturation, the catalytic domain and the C-terminal fibronectin (Fn) III-like domain (Meyer et al., 2016, Nakagawa et al., 2010). In recent years, subtilases have emerged as critical regulators in plant-microbe interactions (Schaller et al., 2018). For example, during immune responses, plants utilize SBT5.2a to regulate the abundance of elicitors in the apoplast, owing to its ability to cleave the 22-residue flagellin epitope (flg22) at multiple sites (Buscaill et al., 2024). Intriguingly, large-scale phylogenetic analysis of plant subtilases, combined with transcriptomic data, has additionally suggested potential roles for subtilases belonging to the SBT1.13 and SBT1.10 lineages in symbiotic interactions (Taylor & Qiu, 2017). Consistent with this, as a representative member from those lineages, the *Casuarina glauca* subtilase gene *Cg12* is specifically induced in response to *Frankia* infections (Fournier et al., 2018, Svistoonoff et al., 2003, Laplaze et al., 2000), while its function remains unsolved. The same applies to the subtilases *SbtM1*,*3*,*4* and *SbtS* that are highly activated during arbuscular mycorrhiza symbiosis (AMS) (Kistner et al., 2005, Takeda et al., 2009, Takeda et al., 2011). However, genetic evidence for the importance of these symbiotic subtilases or of their target proteins have, to our knowledge, never been reported so far. Here, we study a Medicago subtilase, SBT12a, that is specifically induced during RNS and we demonstrate that SBT12a is genetically required for symbiosome maintenance and nitrogen fixation.

## Results

### SBT12 subtilases are symbiotically regulated in legumes

Although the compartmentalization of efficient nitrogen-fixation in symbiosomes is at the heart of RNS, the mechanisms driving symbiosome stabilization and maturation are sparsely understood. To identify core genetic components, we searched available legume transcriptome data for genes with peaking expressions during nodule formation including their nitrogen-fixing state. These data were cross-referenced with evolutionary datasets, pre-assuming that the core gene set should have been retained in nodulating species. Among them, two proteases, Medtr7g079300 and Medtr7g079310 sparked our interest as their expression strongly correlated with leghemoglobin genes and was associated with symbiotic tissues (Extended Data Fig. 1a). Phylogenetically, both genes clustered with members of the subtilase SBT1.13 subfamily (Fig. 1a), the lineage that was potentially implicated in symbiotic associations (Taylor & Qiu, 2017). According to the previous nomenclature within this clade (Taylor & Qiu, 2017), we named these subtilases SBT12a (Medtr7g079300, MtrunA17Chr7g0248861) and SBT12b (Medtr7g079310, MtrunA17Chr7g0248871). Interestingly, they are the only members of this clade being significantly upregulated during RNS, based on the available transcriptome data (Schiessl et al., 2019). (Fig. 1a).

**Fig. 1:**
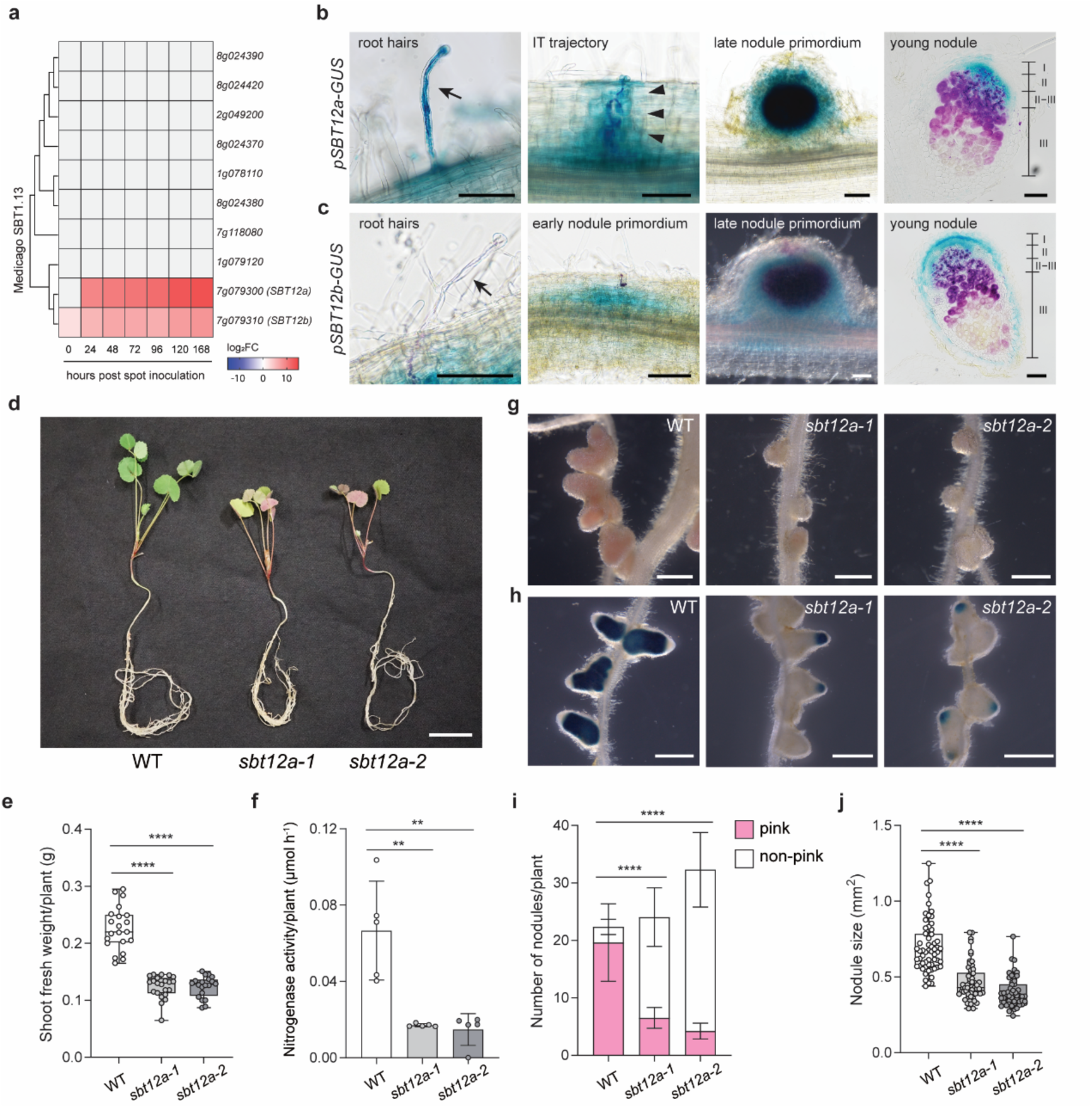
*SBT12a* is induced during rhizobial symbiosis and affects nitrogen fixation. **a**, Phylogenetic and expression analysis of Medicago subtilases belonging to the SBT1.13 clade. The phylogenetic tree was generated from the alignment of subtilase sequences retrieved from the Phytozome v13 database (https://phytozome-next.jgi.doe.gov/) using the neighbor-joining method. The symbiosis-related expression data for each gene were obtained from the previous study (Schiessl et al., 2019). **b**,**c**, Promoter-GUS assays visualizing the expression domains of *SBT12a* (**b**) and *SBT12b* (**c**) throughout the nodulation process. GUS activity and *Sinorhizobium meliloti 2011*-expressed *LacZ* were stained blue and magenta, respectively. Arrows indicate infection threads (ITs) in root hair cells and arrowheads indicate ITs penetrating from epidermis to inner cortical layers (IT trajectory). Different zones were marked in longitudinal sections of elongated nodules. zone I, meristem; zone II, infection zone; zone II-III, interzone; zone III, fixation zone. Scale bars, 100 μm. **d**, *sbt12a* mutant seedlings displayed nitrogen-defective growth phenotype at 21 days post inoculation (dpi) with *S. meliloti 2011*. Scale bar, 2 cm. **e**, Representative nodules formed on WT and *sbt12a* mutant roots at 21 dpi with *S. meliloti 2011*. **f**, WT and *sbt12a* mutant nodules at 28 dpi with *S. meliloti 2011* carrying a *pnifH::GUS* reporter enabling visualization of zones with active nitrogen fixation. Scale bars, 1 mm (**e** and **f**). **g**, Shoot fresh weight per each plant was quantified. n=21, 22 and 20 for WT, *sbt12a-1* and *sbt12a-2*, respectively. Statistically significant differences were determined using Mann-Whitney test. **** p<0.0001. **h**, Acetylene reduction assay (ARA) showing the disrupted nitrogen fixation capacity of *sbt12a* mutant compared to WT plants. n=5 for all genotypes. Each data point denotes the mean value of 5 plants. Error bars represent standard deviation (SD) and statistically significant differences were analyzed using Mann-Whitney test. ** p<0.01. The ARA was independently repeated twice with similar results. **i**, Quantification of nodule number and type in **e**. n=21, 22 and 20 for WT, *sbt12a-1* and *sbt12a-2*, respectively. Error bars represent SD and number of pink nodules formed between WT and mutants were statistically analyzed based on Brown-Forsythe and Welch ANOVA test followed by Dunnett’s multiple T3 comparisons. **** p<0.0001. **j**, Nodule size of WT and *sbt12a* mutants. n=59, 57 and 55 for WT, *sbt12a-1* and *sbt12a-2*, respectively. Kruskal-Wallis test followed by Dunn’s multiple comparisons test was applied for statistical analyses. **** p<0.0001. Phenotyping experiments mentioned above were independently repeated four times with similar results.

We first recapitulated their transcriptional induction by quantitative Real-Time PCR (qRT-PCR) using a cDNA time course covering root infection and nodule development (0-14 days post inoculation, dpi). We confirmed significantly increasing transcript levels over time for both *SBT12a* and *SBT12b* (Extended Data Fig. 1b,c). To assess spatiotemporal expression dynamics over the nodulation process, we cloned their putative promoter regions, selecting 2000 bp upstream of the translational start sites of both *SBT12a* and *SBT12b* to generate beta-glucuronidase (GUS) reporters. These constructs were transformed into Medicago A17 wild-type (WT) roots, which were subsequently inoculated with *Sinorhizobium meliloti* 2011 expressing a beta-galactosidase (lacZ) reporter. Assessing GUS activity at different symbiotic stages, we detected an activation of the *SBT12a* promoter in infected roots hairs, along the IT trajectory in the root cortex, in nodule primordia and predominantly in the infection/interzone (zone II/II-III) of mature nodules (Fig. 1b). By contrast, *SBT12b* promoter activation was closely linked to meristematic tissues, as we detected staining in lateral root primordia and root tips of uninoculated roots, nodule primordia, young nodules and in the apical, meristematic region of mature nodules (Fig. 1c and Extended Data Fig. 1d). These expression patterns are consistent with their spatial transcript abundance retrieved from available single-cell RNA sequencing dataset (Pereira et al., 2024), where *SBT12a* expression was mainly found in putative infected nodule cells that eventually contain nitrogen-fixing symbiosomes, while *SBT12b* transcripts were enriched in tissues with meristematic activity (Extended Data Fig. 1e,f). Thus, both genes are symbiotically induced and expressed during nodulation, but with the clear distinction that *SBT12a* expression is restricted to infection sites while *SBT12b* is more widely associated with dividing cells. Notably, the rhizobial infection-specific expression pattern of *SBT12a* resembles those of its *C. glauca* and *Discaria trinervis* homologs, *Cg12* and *Dt12*, whose expression domains were found to be linked to *Frankia* colonization sites during actinorhizal symbiosis (Svistoonoff et al., 2003, Fournier et al., 2018).

To gain evolutionary insights into the putative importance of *SBT12a* and *SBT12b* during RNS, we reconstructed a phylogenetic tree of *SBT12a*, *SBT12b* and their homologs using 100 eudicot species, including 62 species from the nitrogen-fixing clade (NFC) and 38 plant species outside the NFC as outgroups (Supplementary Table 1), and integrated available transcriptome data into this phylogeny (Libourel et al., 2023). We first uncovered that *SBT12a* and *SBT12b* originate from a recent duplication in the common ancestor of Medicago and *Trifolium pretense,* and belong to a single clade, here named Papilionoid A, whose members specifically respond to RNS (Extended Data Fig. 2a). Notably, *Lotus japonicus*, which forms determinate nodules, carries two *SBT12* homologs, one of which resides in a distinct clade, Papilionoid B, suggesting possible functional divergence associated with nodule type during evolution (Extended Data Fig. 2a). Secondly, by analyzing the presence of *SBT12* genes in those 100 species we found that *SBT12* homologs were frequently lost in non-nodulating species, with few exceptions mainly within the mimosoid clade (Extended Data Fig. 2b). This pattern was particularly striking in Cucurbitales, where *SBT12* homologs were absent in all six non-nodulating species but retained in both nodulating species (Extended Data Fig. 2b). Taken together, these results indicate that *SBT12* homologs are broadly conserved across the NFC, with evidence of functional specialization among certain clade members.

### *sbt12a* mutants are severely altered in nitrogen fixation

To investigate their role in RNS genetically, we identified independent exonic mutant alleles for both genes within the *Tnt1* retrotransposon mutant collection (Tadege et al., 2008). Homozygous individuals were isolated for *sbt12a-1* (NF1441), *sbt12a-2* (NF18072), *sbt12b-1* (NF4378) and *sbt12b-2* (NF19394) (Extended Data Fig. 3a-c). When grown under fully fertilized (high N) conditions, none of the mutant alleles showed any developmental defects upon scoring shoot and root fresh weight and comparing these to WT R108 plants (Extended Data Fig. 3d-i). By contrast, both *sbt12a* mutant alleles developed severe signs of nitrogen starvation (anthocyanin accumulation, chlorotic leaves) when grown in the absence of an inorganic nitrogen source but in the presence of symbiotic *S. meliloti* 2011 at 21 dpi, whereas WT plants remained healthy (Fig. 1d). In line with this, the shoot fresh weight was significantly reduced in the *sbt12a* mutants (Fig. 1e). To verify that nitrogen fixation capacity is altered in the *sbt12a* mutants, we assayed rhizobial nitrogenase activity within the mutant nodules and found that it was almost fully abolished (Fig. 1f). We further examined nodule phenotypes and found that, at 21 dpi, WT plants developed predominantly pink and elongated nodules, whereas both *sbt12a* mutants formed mostly white or greenish nodules, with only a small proportion appearing pale pink and nevertheless less elongated (Fig. 1g,i). Nodules formed on mutant roots were also significantly smaller than those on WT (Fig. 1j). To further validate this symbiotic defect, we inoculated WT, *sbt12a-1* and *sbt12a-2* mutants with *S. meliloti* carrying a *nifH::GUS* reporter that is only active in fixing rhizobia and grew these plants for an extended period of 28 days to allow full nodule elongation. While WT nodules with their extended fixation zone were largely stained, nodules of *sbt12a-1* and *sbt12a-2* exhibited staining restricted to the apical region and displayed an enlarged, non-stained senescence zone (Fig. 1h), consistent with impaired nitrogen fixation. By contrast, none of these phenotypes were observed on *sbt12b* mutant plants (Extended Data Fig. 4a-e).

To determine the developmental basis and gain cellular resolution of the patterns observed in *sbt12a* mutants, we examined nodules across different stages of nodule development (7, 10, 18 and 28 dpi). Toluidine blue-stained nodule sections showed that in WT nodules, rhizobial colonization gradually progressed as expected, with representative nodules displaying characteristic zonation of indeterminate nodule type at 18 dpi (Fig. 2a). A dominant fixation zone with densely packed infected cells and a central vacuole surrounded by numerous bacteria were observed (Fig. 2a,b). In contrast, both *sbt12a-1* and *sbt12a-2* nodules exhibited overall less infected cells at early stages (7 and 10 dpi) compared to WT nodules, indicating a delayed nodulation process in the mutants. Notably, under our experimental conditions, mutant nodules showed signs of premature senescence at 10 dpi, and this phenotype became pronounced at 18 dpi, demonstrated by a markedly enlarged senescence zone (zone IV) containing host cells with degraded contents (Fig. 2c-f). At 28 dpi, WT nodules still retained densely stained infected cells throughout most of the nodule, while *sbt12a* mutant nodules were largely occupied by those senescent tissues (Fig. 2a,c,e). Moreover, approximately a quarter of the *sbt12a* nodules additionally displayed defects in nodule colonization (Extended Data Fig. 5a-e). Semi-thin sections revealed the presence of nITs in the infection zone of WT nodules, which appeared bulky in the *sbt12a-2* mutant. Further ultrastructural analyses using transmission electron microscopy (TEM) confirmed the presence of enlarged nITs with occasional intercellular (apoplastic) bacterial release in the *sbt12a* mutant, whereas these infection structures were mostly confined in WT nodules (Extended Data Fig. 5f,g).

**Fig. 2:**
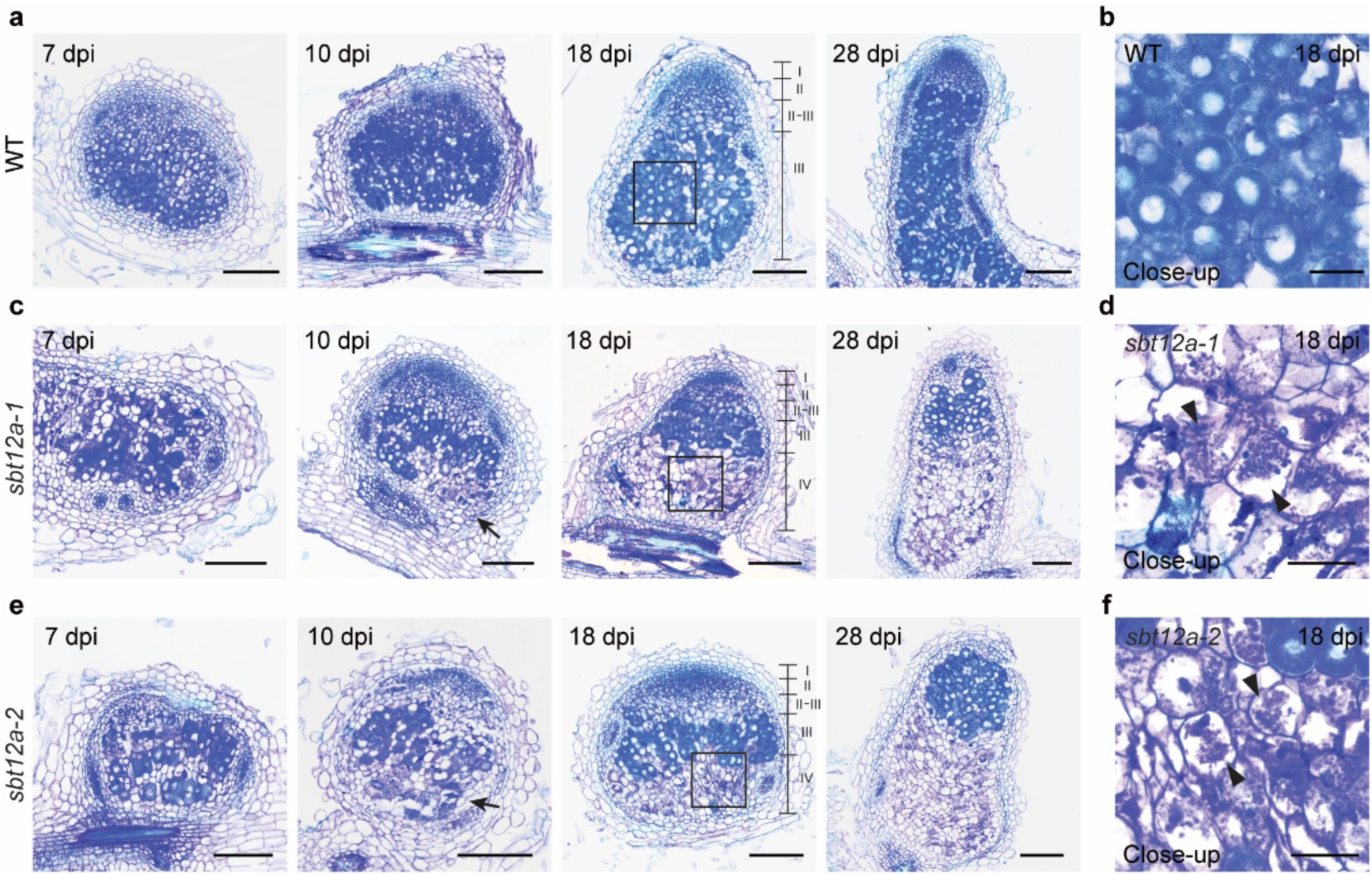
Loss of *SBT12a* function triggers premature nodule senescence. **a**,**c**,**e**, Longitudinal semi-thin sections of nodules from WT (**a**), *sbt12a-1* (**c**) and *sbt12a-2* (**e**) roots inoculated with *S. meliloti 2011*, collected at 7, 10, 18 and 28 dpi. Nodule zones are delineated at 18 dpi. An additional zone IV (senescence zone) is already apparent in *sbt12a* mutant nodules at this stage, indicating premature onset of senescence, whereas the nitrogen fixation zone (zone III) is markedly smaller in *sbt12a* nodules compared to WT. **b**,**d**,**f**, Magnified views of the boxed regions in (**a**,**c**,**e**), respectively. Arrows in the 10-dpi images indicate nodule regions showing signs of senescence. Arrowheads mark individual senescent nodule cells, which are well pronounced in mutant but almost absent in WT nodules. For each time point, nodules used for sections were randomly selected from 10 plants per genotype in two independent experiments. All sections were counterstained with 0.05% (w/v) toluidine blue. Scale bars, 200 μm (**a**,**c**,**e**) and 50 μm (**b**,**d**,**f**).

As both *SBT12a* and *SBT12b* were induced at very early stages of nodulation (Fig. 1a-c and Extended Data Fig. 1a-c), we assessed the rhizobial infection progress and nodule primordium development as early as 7 dpi in WT, *sbt12a* and *sbt12b* mutants. Quantification analyses suggested that the *sbt12a* mutant exhibited slightly delayed infection progression, while *sbt12b* mutants did not significantly differ from WT in any of the event categories examined (Extended Data Fig. 6a,b). From these data we concluded that only *SBT12a*, but not *SBT12b*, actively controls nodule colonization and nitrogen fixation, and therefore only considered SBT12a for the following experiments.

### SBT12a controls bacteroid differentiation in symbiosomes

The premature nodule senescence in *sbt12a* mutants suggested that symbiosome development might be compromised. To test this, we assessed symbiosome and bacteroid states in more detail. TEM analyses of 21 days old nodules revealed that infected cells in the nitrogen-fixation zone of WT nodules contained predominantly elongated, rod-shaped symbiosomes, with a very narrow or barely visible peribacteroid space (Fig. 3a). By contrast, infected cells in *sbt12a* mutant nodules frequently displayed multiple structural defects. In a subset of cells, we observed an increased number of vesicles together with numerous shorter, less elongated symbiosomes, some of which appeared to be fusing with each other (Fig. 3b), possibly indicating the onset of bacterial degradation. In other cells, although the overall cellular architecture was still intact, most bacteroids inside appeared detached from the symbiosome membrane and the peribacteroid space was markedly enlarged (Fig. 3c). Eventually, these infected cells became completely devoid of symbiosomes, leaving only membranous remnants (Fig. 3d). Together, these observations indicate that, in the absence of SBT12a, symbiosomes are not properly maintained, which ultimately leads to early senescence of infected cells.

**Fig. 3:**
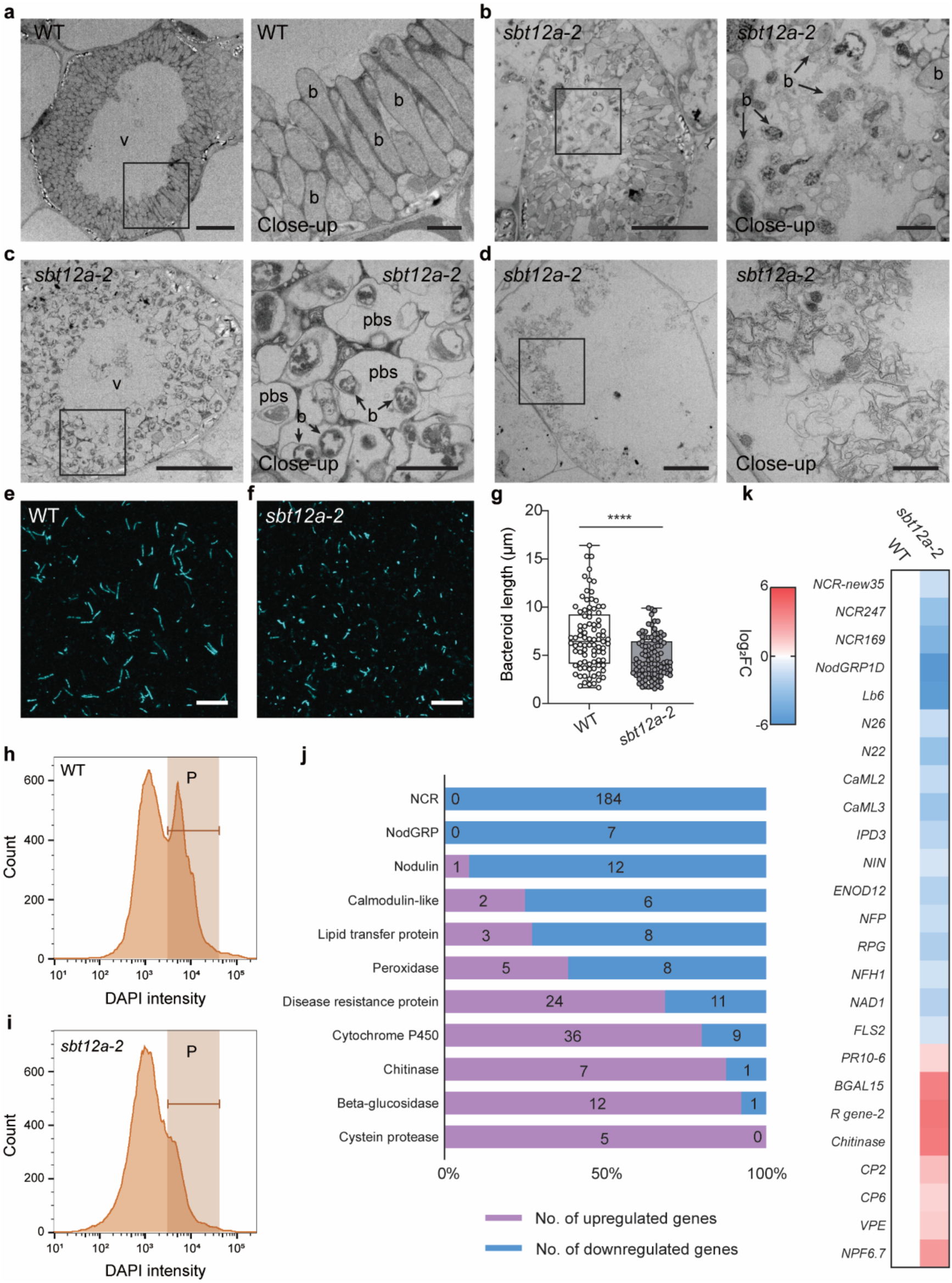
The *sbt12a* mutant is impaired in bacteroid differentiation. **a**-**d**, Transmission electron microscopy (TEM) analysis of infected nodule cells in WT (**a**) and *sbt12a-2* (**b**-**d**). WT host cells are typically filled with elongated bacteroids surrounding a central vacuole (**a**), whereas in *sbt12a-2* mutant nodules increased vesicular events are often observed (**b**), leading to the gradual degradation of symbiosome structures. These symbiosomes are markedly disorganized, with enlarged peribacteroid spaces and multiple small bacteroids enclosed in a single symbiosome (**c**), and ultimately collapse (**d**). b, bacteroid. pbs, peribacteroid space. V, central vacuole. Close-up views of the boxed regions are shown to the right of each panel. Data were collected from ten nodules at 21 dpi for each genotype in two independent replicates. Scale bars, 10 μm (overview images), 2 μm (close-up images). **e**,**f**, Representative confocal images of isolated bacteroids stained with DAPI (cyan) from WT (**e**) and *sbt12a-2* mutant (**f**) nodules collected at 21 dpi. Scale bars, 20 μm. **g**, Quantification of bacteroids length. n=89 and 98 for WT and *sbt12a-2*, respectively. Statistical differences were analyzed using Mann-Whitney test. **** p<0.0001. **h**,**i**, Flow cytometry analysis of DAPI-stained bacteroids isolated from WT and *sbt12a-2* mutant nodules. Up to 50000 bacteroids per sample were examined for each genotype in the analysis. P represents bacteroid populations that exhibit the same DAPI intensity. Experiments for bacteroid phenotyping were repeated three times with similar results. **j**, Classification of representative differentially expressed genes (DEGs) across gene families. Numbers of up-regulated and down-regulated genes (*sbt12a-2* vs WT) are indicated for each gene family, and are labelled with pink and blue backgrounds, respectively. **k**, The expression of representative DEGs from transcriptome data.

To quantify these differences in bacteroid morphology robustly, we isolated bacteroids from WT and *sbt12a* mutant nodules by fractionation (Fig. 3e,f). Indeed, bacteroids from WT nodules were significantly longer than those from *sbt12a-2* nodules (Fig. 3e-g). This indicates that *sbt12a*-derived bacteroids may not reach the differentiation degree and ploidy level observed in WT nodules. As this can be monitored by the DNA content of these cells, we conducted flow cytometry analysis (FCA) on DAPI-stained bacteroids, considering DAPI intensities of around 10^4^ relative fluorescence units (RFU) as being derived from terminally differentiated bacteroids (Gao et al., 2023). The FCA profiles of bacteroids isolated from young WT nodules showed two dominant peaks, indicating a mixture of young differentiating and older, terminally differentiated bacteroids (Fig. 3h). By contrast, the single FCA peak around 10^3^ RFU suggested that bacteroids in the *sbt12a* mutant failed to undergo terminal differentiation (Fig. 3i).

Although some bacterial release does occur in *sbt12a* nodules, most of these symbiosomes never reach a fully functional stage. To gain a global insight into how *SBT12a* contributes to symbiosome functionality, we performed bulk RNA sequencing on WT and *sbt12a-2* nodules. We first analyzed the reads mapping to *SBT12a* transcripts in the *sbt12a-2* mutant allele. Indeed, a clear deletion within the *SBT12a* transcripts was observed immediately downstream of the *Tnt1* retrotransposon insertion site, rendering these transcripts dysfunctional (Extended Data Fig. 7a). Subsequent comparison of the transcriptome profiles revealed the transcriptional induction of 1161 genes in *sbt12a-2*, while 1142 genes were downregulated (Extended Data Fig. 7b and Supplementary Table 2). When assigning functional categories to these genes, we found a strong repression of classical symbiotic markers in the mutant while several senescence markers including *CYSTEINE PROTEASE* (*CP*) *2*, *CP6* (Berrabah et al., 2023) and *VACUOLAR PROCESSING ENZYME* (*VPE*) (Pierre et al., 2014) were upregulated in *sbt12a* nodules (Fig 3j,k and Supplementary Table 2). This is in line with the fact that *sbt12a* mutant nodules displayed an obvious early senescence phenotype. Furthermore, immunity-related genes, such as *CHITINASES*, *PATHOGENESIS-RELATED PROTEIN* (*PR*) *10* and *FLAGELLIN SENSING 2* (*FLS2*), were also significantly induced in *sbt12a* nodules (Fig. 3j,k), indicating that the suppression of defense responses was compromised in *sbt12a-2* nodules. In addition, the induction of up to 184 prominent regulatory peptides of the NCR family was strongly impaired in the *sbt12a* mutant, consistent with the defective bacteroid differentiation observed above (Fig. 3j and Supplementary Table 2). These includes *NCR169*, *NCR247* and *NCR-new35* (Fig. 3k), for which single loss-of-function mutations have been shown to cause severe defects in symbiosome development and nitrogen fixation (Horváth et al., 2015, Sankari et al., 2022, Horváth et al., 2023). Taken together, these data demonstrate that bacterial differentiation in Medicago nodule cells requires *SBT12a*.

### SBT12a is secreted to bacterial release sites and predominantly into the peribacteroid space

To gain insights into the molecular function of SBT12a, we explored the subcellular localization of the SBT12a protein. For this, we introduced eGFP-tagged SBT12a under the control of its endogenous promoter into Medicago roots via hairy root transformation. This construct was shown to successfully complement the *sbt12a* mutant phenotype (Extended Data Fig. 8e). Confocal microscopy analysis revealed that the fluorescent signal was robustly detected in infected cells within transformed nodules (Fig. 4a). Here, the SBT12a-eGFP fusion protein was initially found to specifically accumulate as small foci around IDs within the infection zone (Fig. 4b). Moreover, the signal of SBT12a-eGFP was later predominantly observed in symbiosomes within the fixation zone but was excluded from bacteroids, indicating a peribacteroid space localization pattern of SBT12a (Fig. 4c,d). To rule out that this signal derives from auto-fluorescent molecules within this space, we conducted Fluorescence Lifetime Imaging Microscopy (FLIM) and confirmed lifetimes of 2.58 ns and 1.51 ns for eGFP and mCherry, respectively (Fig. 4e). These lifetimes resembled values reported earlier for these genetically encoded fluorophores (Lace et al., 2023). These results suggest SBT12a to be a secreted protein in nodule cells, consistent with the apoplastic localization reported for most plant subtilases (Schaller et al., 2018). To further validate the association of SBT12a with symbiosomes, we performed subcellular fractionation followed by Western blot analysis using nodules from *sbt12a-2* mutant roots expressing *SBT12a-eGFP*. Consistent with microscopic observations, SBT12a-eGFP was found to be strongly enriched in the symbiosome fraction (Fig. 4f). This confirmed the localization of SBT12a-eGFP in symbiosomes (Fig. 4g), and additionally supports that our isolation protocol is suited to enrich intact symbiosomes from root nodules.

**Fig. 4:**
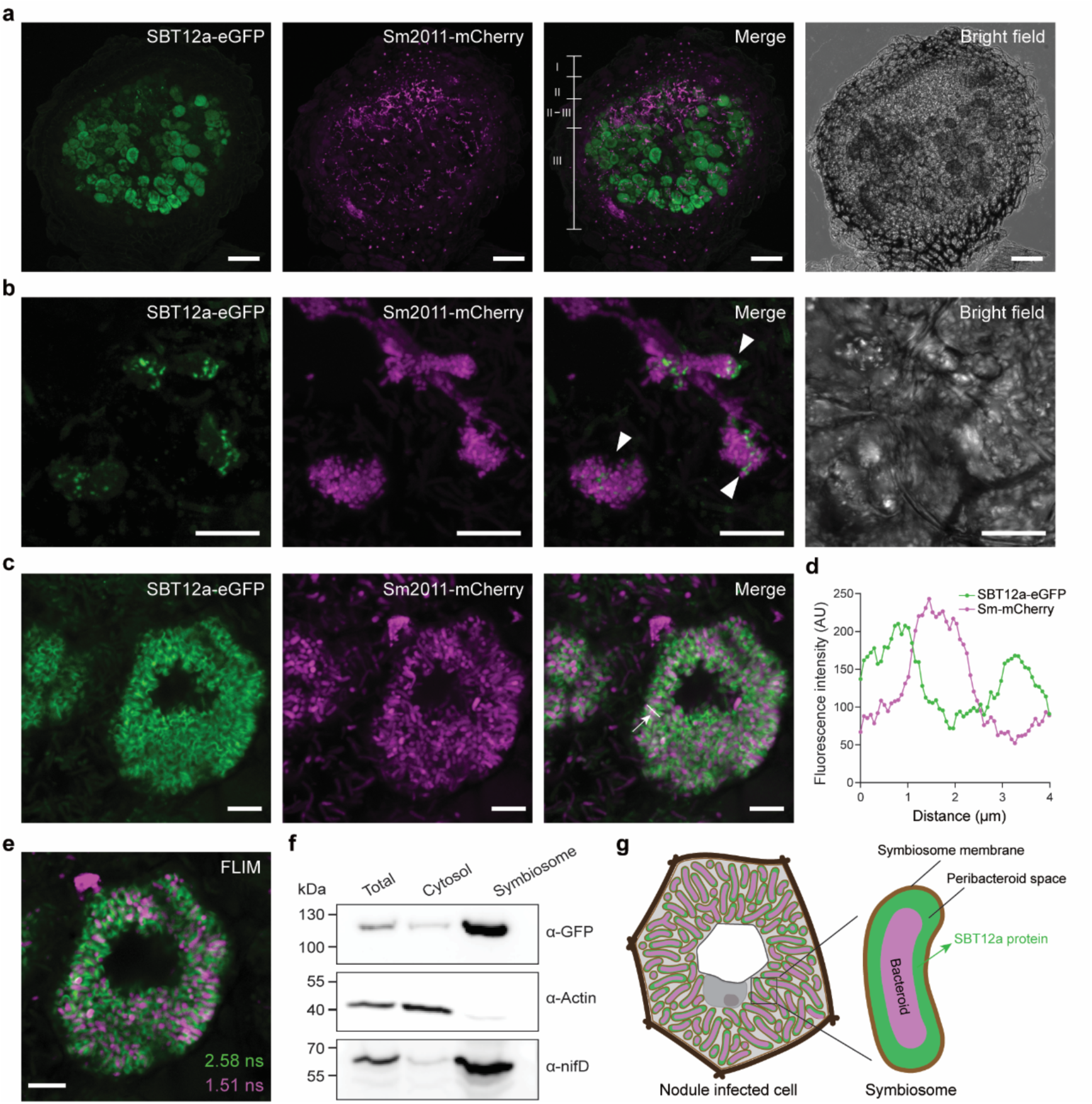
SBT12a localizes at rhizobial release sites and at the peribacteroid space in infected nodule cells. **a**, Confocal microscopy images showing the localization of a SBT12a-eGFP (green) fusion protein in the interzone (zone II-III) and the nitrogen fixation zone (zone III) in nodules. Transgenic nodules expressing *SBT12a-eGFP* driven by its native promoter were infected with mCherry-producing *S. meliloti 2011* (magenta) for 28 days and 70 μm sections were used for confocal imaging. Scale bars, 200 μm. **b**,**c**, Localization of SBT12a-eGFP (green) at putative rhizobial release sites (**b**) and in the peribacteroid space (**c**) of infected nodule cells. White arrowheads indicate the site of rhizobial release. Scale bars, 10 μm (**b** and **c**). **d**, Fluorescence intensities measured along the linear transect (arrow) from (**c**). **e**, Peribacteroid space localization pattern of SBT12a-eGFP analyzed by Fluorescence Lifetime Imaging Microscopy (FLIM). The lifetimes for eGFP and mCherry measured by FLIM are indicated. More than 20 nodules from multiple transgenic roots were analyzed in three independent replicates. Scale bar, 10 μm. **f**, Immunoblot assay showing that SBT12a-eGFP is targeted to the symbiosome. Protein samples were obtained from the following fractions: Total, total nodule extracts; Cytosol, cytosolic fragments of nodule cells; Symbiosome, isolated symbiosomes. α-actin and α-nifD were used as controls for the cytosolic and the symbiosome fraction, respectively. Western blot analysis was repeated twice from two independent rounds of hairy root transformations. **g**, Schematic illustrations of the SBT12a protein (green) localizing to the peribacteroid space surrounding bacteroids (magenta) within symbiosomes.

### SBT12a is a canonical subtilase identified as a phytaspase

At the protein level, SBT12a exhibits a classical plant subtilase topology comprised of an N-terminal signal peptide, an autoinhibitory prodomain, a catalytic domain hosting the catalytic triad (D154, H226, S551) intersected by a protease associated (PA) domain and a C-terminal region that shares similarities to a fibronectin III domain (Extended Data Fig. 8a,b). Notably, an Asp residue was observed at the prodomain junction site, indicating that SBT12a likely functions as a phytaspase, a subtype of subtilases that show stringent Asp specificity for substrate cleavage (Extended Data Fig. 8a, b)(Reichardt et al., 2018). To biochemically characterize SBT12a, we ectopically expressed the WT, catalytically inactive (S551A) and putative propeptide cleavage-resistant (D122A) SBT12a variants, each fused to a triple hemagglutinin (HA) tag, in *Nicotiana benthamiana* leaf epidermal cells. Apoplastic fluids (AFs) were additionally isolated to determine the secretion of SBT12a. As expected, immunoblot analysis revealed two bands for WT SBT12a in total extracts, with the predominant one representing the processed form, which was also present in the apoplast fraction (Extended Data Fig. 8c). In contrast, SBT12a variants carrying the D122A, and especially S551A mutations accumulated as the unprocessed zymogen form and were retained intracellularly (Extended Data Fig. 8c). This observation suggests that SBT12a is autocatalytically processed *in planta* and that its serine activity is required for the zymogen maturation and secretion. We next investigated whether the function of SBT12a depends on its secretion and catalytic activity in nodulation. For this, we generated constructs expressing either WT *SBT12a* or its mutant variants under the control of the native *SBT12a* promoter. Complementation assays showed that only WT *SBT12a* was capable of fully rescuing the nodulation phenotype of *sbt12a* mutants, whereas none of the mutant versions did (Extended Data Fig. 8d,e). Microscopic examination of semi-thin nodule sections further confirmed that complementation of WT SBT12a prevented the induction of premature senescence. By contrast, nodules expressing the SBT12a mutant variants still exhibited this phenotype (Extended Data Fig. 8e). These results indicate that the proteolytic activity is essential for SBT12a to support rhizobial symbiosis.

To further test the putative phytaspase identity of SBT12a and also gain information on its possible target proteins, we first aimed to determine the cleavage specificity of SBT12a. For this purpose, SBT12a was transiently expressed as an HA-fusion protein in *N. benthamiana* leaves and enriched by immunoprecipitation (IP) using HA-traps. The empty vector (EV) and catalytically inactive SBT12a^S551A^ variant were included as negative controls. To assess whether our recombinant SBT12a has serine hydrolase activity, we incubated IP fractions with a Tetramethylrhodamine fluorophosphonate (FP-TAMRA) probe, which specifically and covalently labels serines within enzymatically active serine hydrolases. Indeed, we observed a clear and strong fluorescent signal at around 90 kDa for the WT SBT12a but neither for the SBT12a^S551A^ variant nor the EV control (Fig. 5a). Immunoblotting with an anti-HA antibody confirmed that the SBT12a^S551A^ variant was enriched to a level comparable to WT SBT12a (Fig. 5a), indicating that the lack of labelling reflects loss of catalytic activity rather than reduced protein abundance. These results demonstrate that SBT12a-3xHA enriched by IP is an active serine hydrolase. Next, we analyzed substrate specificity of SBT12a by using a Proteomic Identification of protease Cleavage Sites (PICS) assay (Schilling & Overall, 2008). For this, we incubated our IP-enriched SBT12a protein with an *Escherichia coli* proteome-derived tryptic peptide library and analyzed the corresponding cleavage products by mass spectrometry. This identified 205 cleaved peptides that accumulated after incubation with SBT12a (Fig. 5b and Supplementary Table 3), unambiguously revealing Asp as a dominant residue immediately preceding the cleavage site (P1), and a clear preference for hydrophobic amino acids both upstream (P2 and P3) and downstream (P1’ and P2’) of the scissile bond (Fig. 5c). Moreover, the Asp specificity determined by PICS is consistent with the Asp residue found at the prodomain-peptidase domain junction of SBT12a (Extended Data Fig. 8a,b), as the autolytic processing site generated during maturation generally reflects the substrate specificity of a subtilase (Reichardt et al., 2018). Taken together, our results thus strongly indicated that SBT12a is a *bona fide* phytaspase.

**Fig. 5:**
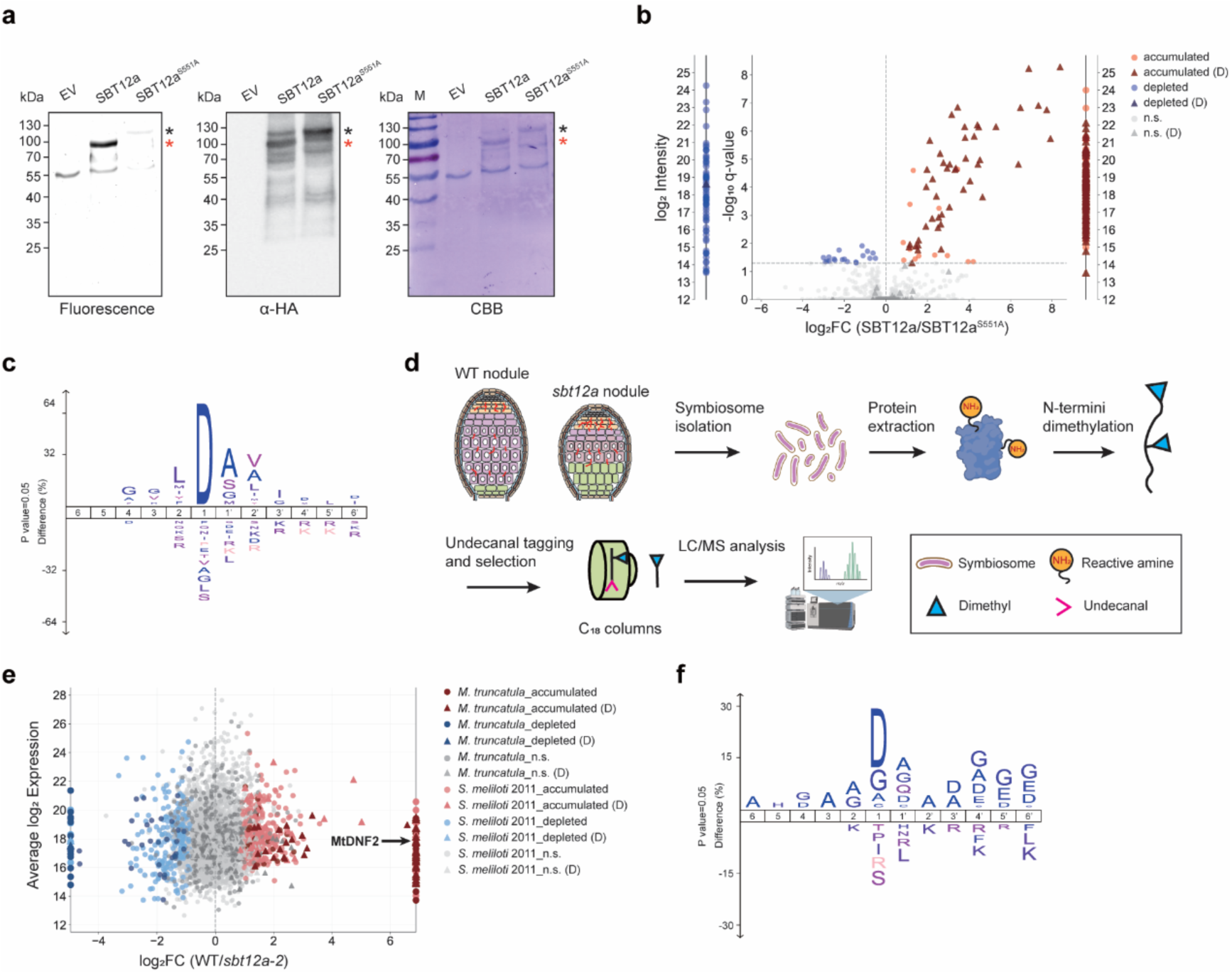
SBT12a mediates Asp-specific proteolytic regulation in symbiosomes. **a**, Immunoprecipitation of empty vector (EV), SBT12a-3xHA, and the catalytically dead mutant SBT12a^S551A^-3xHA (negative control), using *N. benthamiana* leaf extracts. Proteins were labelled with 0.2 μM of the activity-based probe FP-TAMRA and subsequently separated on SDS-PAGE, followed by in-gel fluorescence scanning, immunoblotting with an anti-HA antibody (α-HA) and Coomassie Brilliant Blue (CBB) staining. Black and red asterisks indicate the precursor and mature (self-processed) forms of SBT12a-3xHA, respectively. **b**,**c**, Analysis of substrate specificity of SBT12a by a Proteomics Identification of Cleavage Sites (PICS) assay. (**b**) shows the abundance of peptides quantified in at least two of four biological replicates after incubation with recombinant SBT12a-3xHA or SBT12a^S551A^-3xHA. Peptides detected exclusively in at least three out of four replicates of one condition but absent in the other are displayed on log₂-transformed intensity scales on either side of the volcano plot. Light and dark red indicate increased abundance, light and dark blue indicate decreased abundance, and grey indicates no significant change after treatment with SBT12a compared to SBT12a^S551A^ as determined by a LIMMA-moderated *t*-test with FDR correction (q < 0.05). Triangles mark peptides resulting from cleavage after Asp. (**c**) is the IceLogos showing over- and underrepresented amino acid residues observed preceding (positions 1-6) and following (positions 1’-6’) SBT12a cleavage sites derived from the significantly accumulating peptides. **d**, Schematic representation of the High-efficiency Undecanal-based N Termini EnRichment (HUNTER) pipeline. Some components were created with BioRender.com. **e**, Abundance of identified N-terminal peptides in WT compared to the *sbt12a-2* mutant, with N-termini identified in only one genotype plotted on one side. Peptides shown in red represent N-terminal peptides that were either significantly enriched (q < 0.05) or exclusively found in at least three out of four biological replicates in WT. Blue indicates N-terminal peptides significantly enriched or exclusively observed in *sbt12a-2*. Triangle symbols denote peptides resulting from cleavage with Asp preceding the cleavage site. **f**, IceLogo representations of the amino acid sequence context surrounding the scissile bond, derived from N-terminal peptides that were exclusively and significantly enriched in the WT background.

### N-terminomics of symbiosomes identifies candidate substrates of SBT12a

Having characterized SBT12a as a phytaspase (Fig. 5c) and demonstrated its localization to the peribacteroid space (Fig. 4c-g), we next aimed to identify its in vivo targets within this cellular compartment. To this end, we applied High-efficiency Undecanal-based N-Termini Enrichment (HUNTER), a method for identification of protein N-termini that allows unbiased protease substrate identification (Weng et al., 2019). We isolated intact symbiosomes from WT and the *sbt12a-2* mutant and extracted the corresponding proteins. Primary amines including free amino-termini of in vivo cleaved fragments were modified by reductive dimethylation prior to tryptic digestion. The new amino-termini generated by trypsin were subsequently modified with the hydrophobic aldehyde undecanal, enabling their retention on C18 reverse phase cartridges, leaving only the original protein N-termini in the flow-through (Fig. 5d). Accordingly, high affinity substrates of SBT12a are expected to accumulate in their processed (cleaved) form in WT, while remaining in their unprocessed (uncleaved) form in the *sbt12a* mutant samples.

Mass spectrometric analysis identified a total of 19541 N-terminal peptides, with 1740 originating from Medicago, and 17801 from rhizobia. To identify candidate SBT12a cleavage products, we first limited the dataset to reliably quantified peptides (quantified in ≥ 50% of replicates or exclusively detected in ≥ 3 replicates of one condition) mapping to positions within the protein model (position < 8, to exclude freshly synthesized or aminopeptidase-processed proteins). The resulting dataset contained 509 neo-N-terminal peptides from Medicago, with 31 exclusively found in WT but not in *sbt12a-2* mutant nodules and 37 N-terminal peptides significantly enriched (q ≤ 0.05) in WT (Fig. 5e and Supplementary Table 4). In addition, 24 neo-N-terminal peptides of rhizobial origin were solely detected in the WT and 235 significantly accumulated in WT with respect to the mutant. Further alignment of all these 327 protease-generated N-terminal peptides showed a pronounced selectivity for Asp at the P1 position, along with a preference for small nonpolar residues flanking the putative cleavage sites (Fig. 5f). Collectively, these data revealed SBT12a-dependent candidate substrates derived from both the host plant and its rhizobial symbiont, with potential involvement in RNS.

### SBT12a contributes to the proteolysis of DNF2 in symbiosomes

Among the candidate substrates identified in the N-terminome analysis, Medicago DNF2 attracted our attention based on two reasons. First, peptides mapping to the DNF2 were found to be exclusively present in the WT and harbor Asp at P1 (Fig. 5d and Supplementary Table 4). Second, previous studies have reported an essential role of DNF2 in maintaining bacteroid viability during nitrogen fixation. Moreover, loss-of-function *dnf2* mutant nodules exhibit defense-like responses and premature senescence phenotypes, largely reminiscent of those observed in *sbt12a* mutants.

To test whether DNF2 is indeed a target of SBT12a, we first examined its subcellular localization. As *DNF2* gene transcript was shown to have up to six splice variants in nodules, we specifically focused on the most abundant one, which encodes a 334-amino-acid protein with a predicted N-terminal signal peptide. Constructs for the expression of C-terminal-tagged fusion proteins under the control of the endogenous promoter were generated and introduced into Medicago roots. At 21 dpi, transformed nodules were harvested for analysis. Confocal microscopy revealed that the DNF2-eGFP fusion localized to symbiosomes, presumably the peribacteroid space, in infected cells (Fig. 6a). Nodule fractionation followed by immunoblotting confirmed the presence of DNF2 in symbiosomes, though a signal was also detected in the cytosolic fraction (Fig. 6b). Intriguingly, several bands were enriched in the symbiosome fraction but absent from the cytosolic fraction, suggesting compartment-specific post-translational modifications of DNF2, likely involving proteolytic processing (Fig. 6b).

**Fig. 6:**
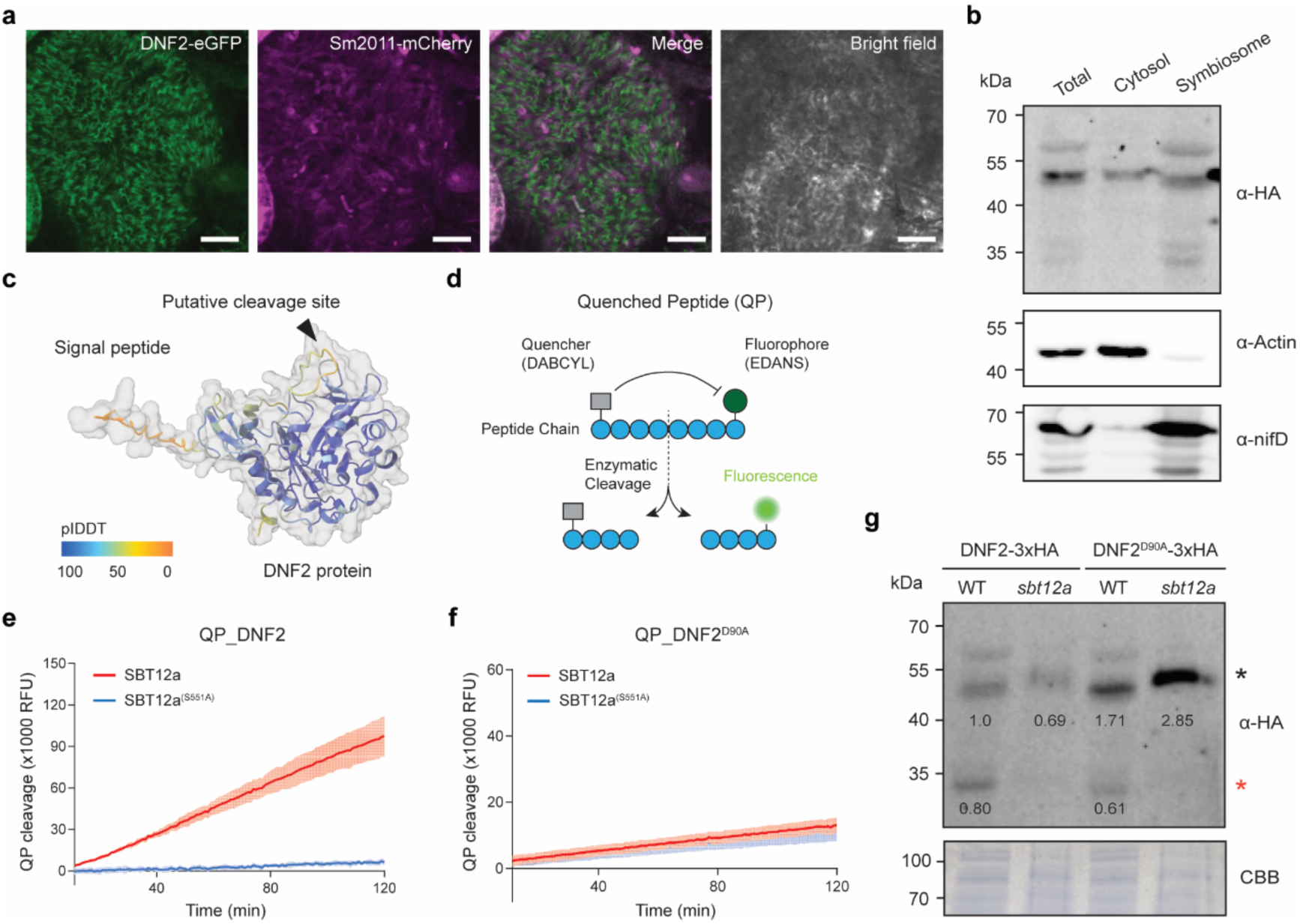
Proteolytic processing of DNF2 within symbiosomes is associated with SBT12a activity. **a**, Confocal microscopy images showing the localization of DNF2-eGFP (green) in a symbiotic cell at 21 dpi with *S. meliloti* 2011-mCherry (magenta), with the DNF2-eGFP signal surrounding individual bacteroids. Scale bars, 10 μm. **b**, Immunoblot assay showing the subcellular fractionation of DNF2-3xHA. Total nodule extracts (Total), cytosolic fragments (Cytosol) and symbiosome fractions (Symbiosome) were isolated from transformed nodules and analyzed by immunoblotting with HA antibody. Anti-Actin and anti-nifD were used as controls for the cytosolic and symbiosome fractions, respectively. **c**, AlphaFold3-predicted structure of DNF2. Colors are based on its pLDDT confidence score. The signal peptide and the putative cleavage site (arrowhead) detected by HUNTER analysis are indicated. **d**, Conceptual illustration of the quenched peptide (QP) assay to monitor processing. **e**, in vitro cleavage of the QP corresponding to the putative cleavage site in DNF2 by recombinant SBT12a-3xHA or the catalytically dead SBT12a^S551A^ mutant, monitored as an increase in relative fluorescence units (RFU) over time. The lines represent the mean value of four independent replicates and the shading represents the SD. **f**, In vitro cleavage assay as in (**e**) using the QP carrying a point mutation (Asp to Ala) which fully abolishes SBT12a-dependent processing. **g**, Immunoblot analysis of symbiosome fractions isolated from nodules formed at 24 dpi on transgenic roots of the WT or the *sbt12a-2* mutant expressing DNF2-3xHA or the putative cleavage-resistant mutant DNF2^D90A^-3xHA, probed with an HA antibody. Similar amounts of protein were loaded into each lane, as shown by the CBB-stained membrane. The processed form of DNF2-3xHA (red asterisk) is absent in the *sbt12a-2* mutant background, while DNF2^D90A^-3xHA is detected predominantly as the full-length protein (black asterisk) in both genotypes, suggesting that DNF2 processing is associated with SBT12a activity in vivo. Numbers indicate integrated intensities that were normalized to the full-length DNF2 band in the left WT lane.

The putative cleavage site identified by our N-terminome analysis locates between position D90 and S91 of DNF2. AlphaFold3 predicted this region to be a flexible, surface-exposed loop between an α-helix and a β-strand, albeit with relatively low confidence (Fig. 6c). Unfortunately, and despite multiple attempts, we were unable to purify correctly folded DNF2 protein in vitro to experimentally test this further. Nonetheless, the putative association between SBT12a and DNF2 was also supported by structural modelling by AlphaFold3, which positioned this cleavage site adjacent to the catalytic Ser of SBT12a (Extended Data Fig. 9a). To directly verify this cleavage event, we adopted a strategy using the quenched peptide (QP) that has been recently applied in other protease assays (Buscaill et al., 2024, Chen et al., 2024). In the intact peptide, close proximity between the quencher and the fluorophore in the non-cleaved peptide prevents fluorescence emission, an effect that is abolished upon peptide cleavage (Fig. 6d). For this, we designed and custom-synthesized the QP containing the WT D90 cleavage site (LSIDSKIS (QP_DNF2)), and exposed it to recombinant, purified, active SBT12a as well as the catalytically inactive SBT12a^S551A^ variant. Indeed, we observed increasing level of fluorescence emission over time when incubating QP_DNF2 with SBT12a but not with the catalytically inactive SBT12a^S551A^ (Fig. 6e). To further test if this cleavage depends on the presence of the central Asp, we replaced Asp with Ala and obtained the corresponding QP (LSIASKIS (QP_DNF2^D90A^)). Incubation of QP_DNF2^D90A^ with SBT12a did not lead to the increase in fluorescence (Fig. 6f), indicating that the cleavage by SBT12a requires the central Asp in the peptide.

Following the results from the QP assay, we investigated whether DNF2 processing requires D90 and depends on SBT12a not only in vitro but also in nodule symbiosomes. For this, a construct to express DNF2-3xHA fusion protein driven by its native promoter was introduced into Medicago WT or *sbt12a-2* mutant roots. We then harvested 24 days old nodules, extracted proteins from isolated symbiosomes and performed immunoblot analysis. In extracts from WT symbiosomes, three bands were detected, with the middle one approximately matching the calculated size of full-length DNF2-3xHA (42 kDa), and the lower molecular weight band matching that of the cleavage product (32 kDa) (Fig. 6g). Image quantification indicated that, in WT samples, the overall abundance of DNF2 full-length protein was comparable to that of the putative cleavage product. By contrast, the abundance of full-length DNF2 was reduced by approximately 30% in the *sbt12a* mutant, while the putative cleavage product was undetectable (Fig. 6g). This suggests that SBT12a contributes to DNF2 processing and may also influence its overall abundance. When expressing the DNF2^D90A^ variant in both WT and *sbt12a* mutant nodules, however, we observed an about two-fold accumulation of full-length DNF2^D90A^, indicating that overall protein stability of this variant is enhanced. For this variant, the full-length protein was about three-fold more abundant than the cleaved form, demonstrating a clear reduction of DNF2^D90A^-processing in the WT. This effect was even stronger in the *sbt12a* mutant, where no cleavage product was detected despite the strong accumulation of the DNF2^D90A^ full-length variant (Fig. 6g). These results support that proteolysis of DNF2 at position D90 in symbiosomes is associated with SBT12a activity. Finally, we aimed to evaluate the functional relevance of this cleavage site within DNF2. For this, we obtained homozygous seeds for a *dnf2* mutant allele (Bourcy et al., 2013) and compared this with WT roots when both being transformed with an empty vector control (Extended Data Fig. 9b). While the WT formed functional pink nodules, these were not observed on *dnf2* mutant roots. Expression of a DNF2-3xHA construct from the endogenous *DNF2* promoter rescued the *dnf2* mutant, as pink nodules were observed, albeit at a lower number than on the WT control. When expressing DNF2^D90A^ in the *dnf2* mutant background, the number of pink nodules was significantly lower compared to roots expressing WT DNF2 (Extended Date Fig. 9c). Additional microscopic inspection of these DNF2^D90A^ nodules revealed often only partial colonization (Extended data Fig. 9b). These data support the role of the DNF2 cleavage mediated by SBT12a.

## Discussion

During legume-rhizobial interactions, symbiosome formation and differentiation represent the key steps in the transition from initial infection and organogenesis to nitrogen fixation within nodules (Zhang et al., 2024). To accommodate thousands of those newly formed organelles in a single cell, the host undergoes extensive cellular adaptations involving the reorientation of endocytic/exocytic pathways (Harrison & Ivanov, 2017), structural reconfigurations of common organelles, in particular vacuoles and the ER (Zhang et al., 2022, Yang et al., 2023), and fine-tuning of various signaling responses (Ren et al., 2025). Intriguingly, recent studies have underpinned the importance of symbiosomes for the evolutionary stabilization of nitrogen fixation in legumes (de Faria et al., 2022, Casaes et al., 2024). However, our knowledge about how host cells tightly monitor and maintain symbiosome integrity over time to achieve sufficient nitrogen fixation is limited. Here, we uncovered the Medicago subtilase SBT12a to be central in this process. Loss of SBT12a impairs symbiotic cell differentiation, leading to a severe defect in nitrogen fixation. Our findings suggest that SBT12a contributes to symbiosome development through the proteolytic regulation of DNF2 and potentially other substrates within the peribacteroid space (Extended Data Fig. 10a,b). Together, our study establishes a critical role for subtilase-mediated proteolysis in sustaining effective RNS.

Expression and phylogenetic analyses identified *SBT12a* and its close paralog, *SBT12b*, as the only two members of the SBT1.13 clade that were transcriptionally induced and highly expressed along nodule development (Fig. 1a). Importantly, our genetic data revealed that only SBT12a is required for symbiotic nitrogen fixation, indicating a functional divergence during the evolution. However, it should be noted that, although *SBT12a* is induced as early as 1 dpi (Extended Data Fig. 1a), its loss causes just mild early phenotypes (Extended Data Fig. 6a,b). This suggests that SBT12a-dependent proteolysis only becomes critical once symbiosomes have formed and begin to differentiate.

Comparable to other canonical plant subtilases (Schaller et al., 2018), SBT12a follows the secretory pathway in a prodomain-dependent manner (Extended Data Fig. 8c). In infected nodule cells, SBT12a is specifically targeted to the peribacteroid space (Fig. 4c-g), a compartment that is topologically equivalent to the apoplast and serves as a unique interface for the interkingdom molecular trafficking required for nitrogen fixation (Zhang et al., 2024). It is therefore tempting to speculate that SBT12a is subjected to nodule specific secretory pathway mediated by the DNF1-containg signal peptidase complex (Wang et al., 2010). This is supported by the fact that both *SBT12a* and *DNF1* shared largely overlapped expression domains in the infection zone of mature nodules (Fig. 1b) (Wang et al., 2010). It is worth noting that the expression domain of *SBT12a* in nodules differs from the localization of the protein, which accumulates throughout the fixation zone (Fig. 4a). This suggests that SBT12a remains stable as symbiosome develop. Such spatial separation has also been shown for other symbiosome (membrane)-localized proteins such as SYMREM1 and NCR peptides (Lefebvre et al., 2010, Horváth et al., 2023).

In line with the presence of a characteristic Asp at the prodomain junction, our PICS assay identified that SBT12a functions as an Asp-specific phytaspase (Fig. 5b,c), with a substrate preference closely resembling those of rice and tomato phytaspases . While phytaspases have so far been implicated in stress-induced cell death and flower abscission (Chichkova et al., 2010, Reichardt et al., 2020), our results reveal a crucial role for a phytaspase in regulating RNS. To identify candidate SBT12a substrates, we purified symbiosomes from nodule cells and subjected them to N-terminomics. This approach reduces background from the host cytosol and focuses on proteolytic events occurring in the compartment where SBT12a is active. HUNTER analysis revealed SBT12a-dependent cleavage events (Fig. 5e,f and Supplementary Table 4), suggesting that SBT12a shapes the N-terminome of symbiosomes. However, we cannot rule out that some of the cleaved products may arise from other proteolytic events rather than direct processing by SBT12a. Importantly, several SBT12a-dependent N-termini identified in vivo by HUNTER match the specific cleavage motif, with a central Asp residue at the P1 position determined in vitro by PICS (Fig. 5c,f). This indicates that these N-termini likely derive from SBT12a activity.

DNF2 emerged as a particularly interesting candidate substrate of SBT12a because of its defined role in maintaining functional nitrogen-fixing symbiosomes (Bourcy et al., 2013). However, despite containing a PI-PLC X domain (PI-PLC-XD), DNF2 lacks part of the catalytic core, and the mechanism by which it dampens defense-like responses during bacteroid differentiation remains unclear (Bourcy et al., 2013). Recent studies have reported phospholipase activity for human PLCXD proteins, such as PLCXD1 and PLCXD2 (Verpoort et al., 2025, Frendo-Cumbo et al., 2026), raising the possibility that DNF2 may also possess lipid-hydrolyzing activity in nodule cells. Another peculiar feature of DNF2 is its predicted N-terminal signal peptide, which directs the protein to the secretory pathway and targets it into the peribacteroid space (Fig. 6a,b). This localization differs from that reported for other plant PLC proteins, which mostly associate with the plasma membrane or cytoplasm (Laxalt et al., 2025), and may reflect a specialized role of DNF2 in rhizobial symbiosis. Notably, our results indicate that DNF2 is post-translationally modified in symbiosomes, given the presence of multiple bands in the symbiosome fraction (Fig. 6b). The altered accumulation of DNF2 proteoforms in symbiosomes of the *sbt12a* mutant supports that DNF2 is subjected to SBT12a-dependent proteolytic regulation (Fig. 6g). This process likely involves the Asp residue at position 90 (D90) as this residue is required for SBT12a-mediated peptide processing in vitro (Fig. 6e,f). Consistently, expression of DNF2^D90A^ resulted in an increased abundance of the full-length DNF2^D90A^ and a reduced abundance of the putative cleavage product, particularly in the *sbt12a-2* background (Fig. 6g). Furthermore, the ability of DNF2^D90A^ to complement the *dnf2* mutant is significantly decreased compared to DNF2^WT^ (Extended Data Fig. 9b,c).

Plant subtilases have been implicated in both general protein turnover and limited proteolysis, in which specific cleavage events regulate protein function (Schaller et al., 2018). Whether SBT12a-mediated processing of DNF2 primarily promotes protein degradation or instead regulates its activity remains an important open question. The phenotypic similarity of *dnf2* and *sbt12a* mutants strongly implies a role of limited proteolysis. The cleavage could thus modulate DNF2 stability, activity and/or associations with other proteins in symbiosomes. In this regard, a recent study has predicted DNF2 to be a central interaction hub for rhizobial proteins (Gao et al., 2026). If these interactions are experimentally validated, SBT12a-mediated cleavage could provide a mechanism for regulating DNF2-associated host-microbe interactions, and therefore influencing symbiosome development.

It should, however, be noted that SBT12a likely processes multiple targets beside DNF2. Two observations support this assumption. First, although both *sbt12a* and *dnf2* mutants show impaired symbiosome integrity, their symbiotic phenotypes are not identical. For instance, approximately half of *dnf2* nodules exhibit brown or dark yellow pigmentation near the basal of region (Extended Data Fig. 9b) (Bourcy et al., 2013), a phenotype not observed in *sbt12a* mutants. Second, besides DNF2, our N-terminome analysis identified a broad range of candidate cleavage products derived from both host and rhizobial proteins, most of which have not yet been functionally characterized. In particular, investigating rhizobia-derived candidates is also of great interest given the association of SBT12a with the symbiosome compartment, where host and bacterial proteins can get into direct contact. Supporting evidence for cross-kingdom proteolytic processing of subtilases has recently been gained for *N. benthamiana* SBT5.2a. In that system, SBT5.2a fine-tunes immune responses by directly inactivating the immunogenicity of bacteria-derived molecules, such as flagellin and cold-shock proteins, in the apoplast (Buscaill et al., 2024, Chen et al., 2024). Whether SBT12a similarly targets rhizobial proteins within symbiosomes remains to be tested. Functional validation of these candidate substrates will help clarify how SBT12a coordinates host and bacterial proteolysis during symbiosome maintenance.

In summary, our study identifies SBT12a as a key regulator of nitrogen-fixing symbiosis in Medicago and provides evidence that it contributes to the proteolytic regulation of DNF2. We propose that SBT12a-mediated proteolysis within the symbiosome compartment is part of a broader regulatory mechanism that supports bacterial differentiation and sustained nitrogen fixation.

## Methods

### Plant materials and growth conditions

Two *Medicago truncatula* (Medicago) ecotypes, Jemalong A17, Jemalong *dnf2* (Bourcy et al., 2013) and R108 were used in this study. All *Tnt1* retrotransposon insertion lines including *sbt12a-1* (NF1441), *sbt12a-2* (NF18072), *sbt12b-1* (NF4378) and *sbt12b-2* (NF19394) generated in the R108 background were obtained from Medicago Mutant Database (Oklahoma State University, USA). Medicago seeds were scarified in 96% sulfuric acid (H_2_SO_4_) for ten minutes followed by rinsing six times with sterile tap water. The washing steps were repeated again after treatment with a bleach solution containing 1.2% sodium hypochlorite (NaClO) and 0.1% sodium dodecyl sulfate (SDS) for 1 minute. Seeds were germinated on 1% water agar plates in the dark at 22°C for 20 hours after vernalization at 4°C for 5 days. Germinated seedings were either transferred to pots filled with 1:1 (v/v) mixtures of quartz sand and vermiculite and kept in a growth chamber with 16 hours light/8 hours dark photoperiod at 24°C for one week before rhizobial inoculation (for phenotyping), or subjected to hairy root transformation.

Wild type *Nicotiana benthamiana* plants were grown in a growth chamber (22°C, 16 hours light/8 hours dark).

### Hairy root transformation and rhizobial inoculation

Hairy root transformation of Medicago plants was performed as described previously (Boisson-Dernier et al., 2001). In brief, seed coats were removed from germinated seedlings and a cut was made 5 mm below the cotyledons using a sterile blade. The wound area was then dipped into solid culture of *Agrobacterium rhizogenes* ARqua1 carrying the corresponding binary vector. Transformed seedlings were placed on Fahräeus medium supplemented with 0.5 mM NH₄NO₃ and kept vertically at 21°C in the dark for 3 days, followed by 4 days under a 16 hours light/8 hours dark photoperiod in a growth chamber, with the root part shaded in the darkness. Next, seedlings were transferred into fresh Fahräeus medium and grown for another ten days under the same growth conditions. Positive roots were selected based on nuclear fluorescent selection maker using a stereo microscope. The transformed plants were then transferred to a mixture of quartz sand and vermiculite (1:1) in pots and inoculated with rhizobia after one week.

*Sinorhizobium meliloti* strain 2011 (Sm2011), Sm2011 *hemA::lacZ*, Sm2011 *nifH::GUS* or Sm2011 mCherry were used for the inoculation of Medicago roots. For this, the Sm2011 strain was first cultured on solid TY medium supplemented with appropriate antibiotics for 3 days at 28°C. Freshly grown Sm2011 cells were then used to inoculate liquid TY medium and cultured for 24 hours at 28°C with shaking. Rhizobial cells were harvested by centrifugation at 3000 rpm for 10 minutes, and the resulting pellet was resuspended in liquid Fahräeus medium to a final concentration of OD_600_ = 0.005. Each plant was inoculated with 10 mL of rhizobial suspension. Plants were watered with sterile tap water (30 mL/pot) and fertilized with liquid Fahräeus medium (30 mL/pot) once a week.

### Phylogenetic analysis

To identify putative SBT genes in Medicago, the S8 domain of known SBT proteins was used as query sequences to perform BLAST searches against Medicago genome (https://phytozome-next.jgi.doe.gov/info/Mtruncatula_Mt4_0v1). The obtained SBT sequences were further manually examined by the criteria of the presence of I9 prodomain and all four conserved residues (Asp, His, Asn and Ser) essential for catalytic function in the S8 domain. All SBT sequences were aligned using the ClustalW program with default parameters and a phylogenetic tree was constructed using the neighbor-joining method in MEGA11 software.

To reconstruct the phylogeny of plant subtilases, we first made a database of proteomes from 100 eudicot species, including 29 species from Fabales, 15 species from Rosales, 9 species from Fagales, and 9 species from Cucurbitales (Supplementary Table 1). The sequences of SBT12a (Medtr7g079300) and SBT12b (Medtr7g079310) were aligned against this database using blastp 2.16.0+ (Camacho et al., 2009), with the following parameters: “-outfmt 6 - evalue 10e-5-word_size -max_target_seqs 999999”. The 7570 sequences from this blast search were aligned using MAFFT v7.526 (Katoh & Standley, 2013) with default parameters and trimmed with trimAL v1.4.rev15 (Capella-Gutiérrez et al., 2009) with the parameter -gt 0.05. A preliminary phylogeny was reconstructed using fasttree v2.1.11 using default parameters (Price et al., 2010). This dataset was reduced in multiple steps by keeping only the closest related sequences to SBT12a and SBT12b, to eventually have a dataset consisting of subtilases that originated from a single copy in the common ancestor of land plants. This set of 262 sequences were aligned with MAFFT using the L-INS-I method, followed by trimming with trimAL with the parameter -gt 0.2. Model testing was performed using IQ-tree v2.2.2.3 with the parameter -m TESTNEWONLY (Minh et al., 2020, Kalyaanamoorthy et al., 2017). The best model based on the Bayesian Information Criterion (BIC) was Q.plant+I+R8 (Minh et al., 2021). A more complex model test was done based on this model by additionally including the profile mixture models C10-C60 (Quang le et al., 2008). The best model based on BIC was again chosen, and the tree was reconstructed with that model with IQ-TREE using the following parameters: “-m Q.plant+C60+F+I+R8 --ufboot 1000 --alrt 1000 --mix-opt --wbtl” (Guindon et al., 2010, Minh et al., 2013). Expression levels of genes were determined using the summarized data provided by a previous study (Libourel et al., 2023). For nodulation, up-and downregulation was based on the columns ending on “WSR_Nod_Up” and “WSR_Nod_Down”. Expression during arbuscular myccorhizal symbiosis was extracted for *M. truncatula* from the column starting with “Medtru_Myct” and for *L. japonicus* from the columns “Lotjap_Myc_Down” and Lotjap_Myc_Up. The phylogeny and related data were visualized in R v4.2.2 (Team, 2022) using the tidyverse metapackage and the packages ggtree v3.15.0, ggtreeExtra v1.8.1, ggnewscale 0.5.1, phytools 2.4-4 and ape 5.8-1 (Yu et al., 2016, Revell, 2024, Paradis & Schliep, 2019).

### Molecular cloning

For plasmids expressed in Medicago, the Golden Gate cloning system was used (Weber et al., 2011). The open reading frame of *SBT12a* was amplified from Medicago Jemalong A17 by PCR using a primer pair with the appropriate overhangs designed for subsequent Golden Gate assembly, and cloned into pJET1.2/blunt vector (Thermo Scientific) followed by Sanger sequencing confirmation. The pJET1.2-SBT12a was further used as a template to generate point mutations by PCR in order to eliminate internal BsaI or BpiI restriction sites, or to create active site mutants of *SBT12a* if needed. Putative promoters of both *SBT12a* and *SBT12b* comprising 2 kb upstream of the start codon were retrieved from the Phytozome database (https://phytozome-next.jgi.doe.gov/), and synthesized by Life Technologies after in silico removal of internal BsaI or BpiI restriction sites present in the sequences. For expressing *DNF2* gene in Medicago, the 4 kb promoter and the coding sequence were used (Bourcy et al., 2013). All other functional Level 0 modules used for Golden Gate assembly in this study were obtained from the ENSA project collection (https://www.ensa.ac.uk/). For expression of *SBT12a* and its mutant variants in *N. benthamiana*, the expression vector pPPO70v1HA was used (Li et al., 2022). *SBT12a* and its mutant variants fused to triple hemagglutinin (HA) tag were amplified using the primers containing *SmaI* and *XbaI* restriction sites. The fragments were then ligated into pPPO70v1HA cut with *SmaI* and *XbaI*. All primers used in this study are listed in the Supplementary Table 5. Details of all constructs are listed in the Supplementary Table 6.

### RNA extraction and quantitative Real-Time PCR

Total RNA was isolated from root or nodule samples harvested at indicated time points using the Spectrum Plant Total RNA Kit (Sigma-Aldrich) according to the manufacturer’s instructions. Subsequently, 1 μg of total RNA was used for cDNA synthesis using SuperScript III Reverse Transcriptase (Invitrogen). The resulting cDNA was diluted 1:5 and 1 μL was used for quantitative Real-Time PCR (qRT-PCR) analysis, which was performed using SYBR Green Master Mix (Applied Biosystems) and a StepOnePlus Real-Time PCR System. Gene expression levels were normalized to a ubiquitin gene used as the internal reference (Liang et al., 2021). Primers used for the analysis are listed in the Supplementary Table 5.

### Acetylene reduction assay

Acetylene reduction assay was conducted as described previously (Yun et al., 2023). In brief, nodulated roots of each plant were collected in a 50 mL vial with a flanged rubber septum. Air was replaced with an equal volume of acetylene using a syringe. Samples were then incubated for 2 hours at 28°C before quantification of ethylene by gas chromatography (GC-4100; EQAI, China). The nitrogenase activity was calculated as the amount of ethylene produced per plant per hour.

### Nodule sections

For semi-thin sections, nodules collected at indicated time points were firstly fixed in buffer Z (0.1 M phosphate buffer (pH 7.0), 10 mM KCl, 1 mM MgCl_2_) containing 2% glutaraldehyde at room temperature for 1 hour. After 3 times washing in buffer Z, fixed nodule tissues were dehydrated in a graded ethanol/water series (50%, 70%, 80%, 90% and 100%) for 1 hour each at room temperature and then subjected to Technovit 7100 (Heraeus Kulzer) resin embedding system according to the supplier’s instructions. Longitudinal sections (10 μm thickness) were made using a RM2245 microtome (Leica Biosystems, Germany).

To assess the protein localization of SBT12a, transformed nodules generated via hairy root transformation were freshly collected and embedded in 6.5 % low melting agarose (Biozym Scientific, Hessisch Oldendorf, Germany). Nodule sections (70 μm thickness) were obtained using a VT1000S vibratome (Leica Biosystems, Germany).

### Histochemical staining

GUS staining was performed as reported previously with minor modifications (Lee et al., 2024). Briefly, roots or nodules harvested at various time points were washed thoroughly and fixed in 2% (v/v) glutaraldehyde at room temperature for 1 hour. After rinsing three times in washing solution containing 50 mM phosphate buffer (pH 7.2), the fixed tissues were immersed in GUS reaction buffer, vacuum-infiltrated for 20 minutes, and incubated at 37°C for 2-4 hours. The GUS reaction was terminated by washing the samples in the same buffer. Stained samples were stored in 70% ethanol until imaging or further subjected to X-Gal staining.

For X-Gal staining, nodulated tissues were immersed in buffer Z supplemented with 0.8 mg/mL X-Gal and 5 mM each of K_3_Fe(CN)_6_ and K_4_Fe(CN)_6_. Samples were vacuum-infiltrated for 20 minutes and subsequently incubated at 28°C overnight until sufficient staining was achieved.

Toluidine blue staining of nodule sections was performed as previously described (Xiao et al., 2014).

### Sample preparation for transmission electron microscopy

WT and *sbt12a-2* mutant nodules harvested at indicated timepoints were cut in half and immediately submerged in 4% paraformaldehyde (PFA) and 2.5% glutaraldehyde in MTSB buffer. Vacuum was applied for 15 minutes and samples were left in the fixative for 3 hours at room temperature, transferred to 4°C and kept overnight. After primary fixation, samples were washed five times with MTSB buffer and post-fixed in aqueous 1% O_S_O_4_ for 2 hours at room temperature, washed and *in bloc* stained for 2 hours in 2% UrAc. Dehydration was carried out in a graded ethanol/water series (50%, 70%, 80% and 95%-15 minutes each), followed by 30 minutes in 100% ethanol and 100% acetone. Samples were then embedded in Epoxy resin. Ultrathin sections (70 nm) were obtained using a Reichert-Jung Ultracut-E microtome and examined using a Hitachi 7800 TEM coupled to a Xarosa CMOS camera (Emsis).

### Confocal microscopy and Fluorescence Lifetime Imaging Microscopy (FLIM)

Fluorescence imaging was conducted using a TCS SP8 confocal microscope (Leica Microsystems, Mannheim, Germany). Images were acquired with a 20×/0.75 and a 40×/1.10 (HC PL APO CS2) water immersion lens. A White Light Laser (WLL) was used as an excitation source. eGFP was excited at 488 nm, and the emission was collected at 500–550 nm. mCherry was excited at 561 nm, and the emission was detected in the 600–650 nm range. FLIM analysis was performed as previously described (Lace et al., 2023). All images were processed and analyzed using Fiji/ImageJ software.

### Nodule fractionation and symbiosome isolation

Nodule fractionation and symbiosome isolation was carried out following the procedure described previously (Catalano et al., 2004). In brief, WT and mutant nodules were cut off from roots and homogenized in a prechilled extraction solution (0.5 M sucrose, 50 mM Tris/HCl (pH 7.4), 10 mM DTT and 1 x Protease Inhibitor Cocktail (Rochel)). The homogenate was filtered through two layers of Miracloth and labelled as nodule total proteins. This resulting suspension was further spun at 10000 g for 1 minute at 4°C, and the supernatant and pellet were labelled as cytosol and symbiosome proteins, respectively. The pellet was resuspended in extraction buffer and washed two more times sequentially in 1.5 M and 1 M sucrose buffer each containing 50 mM Tris/HCl (pH 7.4), 10 mM DTT and 1 x Protease Inhibitor Cocktail. The final symbiosome pellet was resuspended in extraction solution for further analysis.

### Flow cytometry assay

To investigate the role of SBT12a in bacteroid differentiation, bacteroids were isolated from WT and *sbt12a-2* mutant nodules as previously described (Horváth et al., 2023), and subsequently stained with 5 μg/mL DAPI. Flow cytometry analysis was performed using a BD LSRFortessa™ flow cytometer (BD Biosciences). A total of 50,000 bacteroids were analyzed, and the resulting data were processed using FlowJo software.

### RNA sequencing analysis

Three biological replicates using WT and *sbt12a-2* mutant nodules harvested at 18 dpi were prepared for RNA sequencing. Total RNA was extracted as described above, and its quality and concentration were assessed using an Agilent 2100 Bioanalyzer. Library preparation and sequencing were carried out by Novogene Co., Ltd. (Beijing, China). Sequencing data were analyzed using the Galaxy bioinformatics platform (https://usegalaxy.eu/) following standard procedures. Briefly, raw reads were quality-checked and mapped to the Medicago reference genome (https://medicago.toulouse.inra.fr/MtrunA17r5.0-ANR/) using the STAR aligner. Gene-level read counts were generated using STAR, and differentially expressed genes (DEGs) were identified with the DESeq2 package, applying a threshold of log_2_ fold change > 1 and adjusted *P*-value < 0.05.

### Transient expression in N. benthamiana and protein extraction

*Agrobacterium tumefaciens* strain GV3101 was used for agroinfiltration of 4 to 5-week-old *N. benthamiana* leaves. Freshly cultivated agrobacteria harboring expression constructs and p19 silencing suppressor were harvested by centrifugation (4000 g, 10 minutes) and resuspended in an infiltration buffer (10 mM MgCl_2_, 10 mM MES (pH 5.6), 150 μM acetosyringone). The cultures were diluted and mixed to obtain a final OD_600_ of 0.7 each. After 2 hours of incubation, bacterial cells were infiltrated into *N. benthamiana* leaves using needle-less syringes. The infiltrated plants were kept in a growth chamber until use.

For total protein extraction, leaves harvested 4 days after infiltration were ground in liquid nitrogen and suspended in extraction buffer containing 50 mM Tris/HCl (pH 7.5), 100 mM NaCl, 0.5% Triton X-100, 10 mM β-mercaptoethanol and 1 x Protease Inhibitor Cocktail (Roche). Extracts were then centrifuged at 16000 g for 10 minutes at 4°C. The resulting supernatant was used for measurement of protein concentrations by the dye-binding Bradford assay (Bio-Rad). Protein extracts were flash-frozen in liquid nitrogen and stored at -80°C until further analyses.

### Isolation of apoplastic fluids

Apoplastic fluids (AFs) were collected as previously reported (Reichardt et al., 2018). Briefly, agroinfiltrated *N. benthamiana* leaves were submerged in ice-cold extraction buffer (50 mM NaH_2_PO_4_/Na_2_HPO_4_ (pH 6.5), 200 mM KCl) and vacuum infiltrated for 5 minutes. Excess water on the surface of leaves was dried off with absorbing paper, and leaves were mounted into a 25 mL syringe placed in a 50 mL tube. AFs were collected by centrifugation at 1500 g at 4 °C for 7 minutes, and stored at -20°C for further use.

### Western blot

For immunodetection, protein samples were separated by 12% SDS-polyacrylamide gel electrophoresis (PAGE), and transferred to a polyvinylidene difluoride (PVDF) membrane (0.2 µm pore size) using the Trans-Blot® Turbo™ semi-dry transfer system (Bio-Rad). After being blocked with 5% (w/v) powder milk diluted in TBS-T for 1 hour at room temperature, the membrane was hybridized with a primary antibody anti-GFP (Takara Bio Cat# 632381; 1:5000), peroxidase-conjugated anti-HA (Roche, Cat# 12013819001; 1:4000), anti-Actin (Sigma-Aldrich Cat# A0480; 1:5000) or anti-nifD ((Ott et al., 2005); 1:5000), at 4 °C overnight. The membrane was washed 3 times in TBS-T for 30 minutes. Peroxidase-conjugated secondary antibodies (anti-mouse, Sigma-Aldrich Cat# A4416; 1:5000) was applied, if required, for 2 hours at room temperature. Signals were visualized using the Clarity Western ECL Substrate (Bio-Rad) on an ECL ChemoCam Imager (Intas Science imaging).

### Immunoprecipitation

Total protein was extracted from *N. benthamiana* leaves transiently expressing the empty vector control, SBT12a-3xHA or SBT12a^S551A^-3xHA using the same extraction buffer as mentioned above but without adding protease inhibitors. 10 mL of each protein sample was incubated with 40 µL HA-Trap Magnetic Agarose (ChromoTek) at 4 °C for 1 hour with gentle rotation. Beads were then washed five times with washing buffer (10 mM Tris/HCl (pH 7.5), 150 mM NaCl and 0.5 mM EDTA). Bound proteins were subsequently eluted using 500 μM HA peptide (ChromoTek), and the protein concentration was determined using the Bradford assay.

### Activity-based protein profiling (ABPP)

FP-TAMRA probe (Thermo Scientific) was used to label and detect active subtilases. The probe was dissolved in 100 μL of DMSO to make a 0.1 mM stock solution. 10 μL of immunoprecipitated SBT12a-3xHA or SBT12a^S551A^-3xHA were incubated with 0.2 μM FP-TAMRA in PBS (pH 7.0) for 1 hour at room temperature in the dark. The reaction was stopped by adding 2 x SDS sample buffer and boiling for 5 minutes. Labeled proteins were analyzed by SDS-PAGE followed by fluorescent gel scanning with a Typhoon FLA 9000 (GE Healthcare Life Sciences) using Cy3 settings (532 nm excitation, 610PB filter).

### Proteomic Identification of Cleavage Sites

An *Escherichia coli* (strain K12) proteome-derived peptide library was generated by whole cell proteome digestion with trypsin as described (Demir et al., 2022a). Aliquots of 5 µg purified tryptic peptide libraries were equilibrated in PBS (pH 7.0) buffer. IP-enriched recombinant SBT12a or catalytically inactive SBT12a^S551A^ were added and incubated for 2 hours at 25°C. Four replicate reactions were performed for each construct. As a modification to the original protocol (Demir et al., 2022b), the resulting peptide samples were not modified by dimethylation but directly stopped by acidification, individually desalted using self-packed three-layer SDB-RPS StageTips, reconstituted in 0.1% TFA and loaded onto Evotips (Evosep) for label-free analysis.

### Enrichment of N-terminal peptides

N-terminal peptides were enriched with the HUNTER method as previously described (Stockert et al., 2024). Briefly, proteomes were extracted with a denaturing extraction buffer (6 M guanidine hydrochloride, 100 mM HEPES (pH 7.4)), followed by reduction with 10 mM DTT and cysteine carbamidomethylation with 50 mM CAA. An estimated 100 μg of protein per replicate was purified with paramagnetic SP3 beads (Cytiva). Primary amines were labelled by reductive dimethylation with heavy formaldehyde (^12^CD_2_O). The reaction was quenched and samples were digested overnight with trypsin (1:100 w/w) at 37°C. Subsequently, 10 μg of peptides were withdrawn to determine labelling efficiency (pre-HUNTER). Newly generated tryptic peptides bearing primary amines were modified with 100 mM undecanal and removed using reversed-phase HRX spin cartridges (Macherey & Nagel). N-terminal peptides remaining in the flow-through were desalted by self-packed three-layer C18 StageTips (Rappsilber et al., 2007) and reconstituted in 0.1% TFA.

### Mass spectrometry data acquisition

Peptide samples were measured on an Exploris 480 mass spectrometer (Thermo Scientific) coupled to nanoflow chromatography systems. For HUNTER samples, peptides were separated with an Ultimate 3000 RSLCnano system (Thermo Scientific), loaded on a μPAC C18 trapping column (Thermo Scientific) with flow rate of 10 μL/minute and separated on a μPAC C18 pillar array analytical column (50 cm bed length; Thermo Scientific) at a flow of 0.5 μL/minute. Both columns were kept at 40°C. Peptides were eluted using a binary solvent system consisting of 0.1% formic acid (solvent A) and 86% acetonitrile with 0.1% formic acid (solvent B). Mobile phase B increased as follows: 1% to 4% over 2 minutes, 4% to 25% over 24 minutes, 25% to 44% over 12 minutes, and a final column wash at 90% B for 4 minutes. Peptide were introduced into the mass spectrometer with a NanoSpray Flex ion source (Thermo Scientific) equipped with the μPAC Flex iON Connect and a fused silica emitter (20 μm inner diameter, 10 μm tip inner diameter, 360 μm outer diameter). The source voltage was set to +1.8 kV and the ion transfer tube temperature to 275°C. Peptides of PICS samples were separated with 44 minutes gradient on an Evosep One system (EvoSep) equipped with a nano-LC column (EV1137, 15 cm × 150 μm, Evosep) maintained at 40°C by a column oven (PRSVO-V2, Sonation). The binary solvent system consisted of 0.1% formic acid (solvent A) and 100% acetonitrile with 0.1% formic acid (solvent B) and operated in the 30 SPD mode. For electrospray ionization, a stainless-steel emitter with an integrated liquid junction (EV1072, Evosep) was mounted into an EasySpray adapter and installed in a NanoSpray Flex ion source (Thermo Scientific). The spray voltage was set to +2.0 kV and the ion transfer tube temperature to 275°C. Data were acquired in data-independent acquisition (DIA) mode. Each acquisition cycle comprised one MS1 survey scan (m/z 350–1400, resolution 120,000, RF lens 40%, AGC target 300%, maximum injection time 45 ms), followed by MS2 fragment ion scans in profile mode (resolution 30,000, RF lens 50%, AGC target 1000%, maximum injection time 54 ms, variable isolation window width with 1 m/z overlap).

### Mass spectrometry data analysis

Acquired DIA data were analysed with FragPipe v24.0 (Yu et al., 2023) with the integrated MSFragger (v4.4.1) search engine. For HUNTER data, peptide spectra were predicted from the Medicago and *S. meliloti 2011* proteome databases, with enzyme specificity set to semi-specific cleavage with variable N-terminus and C-terminal ArgC digest, up to two missed cleavages, peptide length of 7-45 amino acids, precursor mass range of 200-5,000 Da, and a maximum of five variable modifications per peptide. Dimethylation at protein N-termini and lysine side chains (+32.0564 Da) and cysteine carbamidomethylation (+57.0215 Da) were set as fixed modifications. Variable modifications included methionine oxidation (+15.9949 Da), acetylation of peptide N-termini and lysines (+9.9541 Da), N-terminal pyroglutamate formation from glutamine (-49.0829 Da) or glutamic acid (-50.0699 Da), and proline monomethylation (-16.0282 Da). For PICS data, spectra were matched against the UniProt canonical *Escherichia coli* (strain K12) proteome (downloaded 2026-03-02, 4,531 entries), with cleavage specificity set to semi-specific cleavage with variable N-terminus and C-terminal trypsin digest, up to two missed cleavages, peptide length of 7–40 amino acids, precursor mass range of 300–5,000 Da, and a maximum of four variable modifications per peptide. Cysteine carbamidomethylation (+57.0215 Da) was set as fixed modification. Variable modifications included methionine oxidation (+15.9949 Da), and acetylation of peptide N-termini and lysines (+42.01056). Peptide and protein identifications were subjected to stringent quality filtering, with a 1% false discovery rate threshold applied at both the peptide-spectrum match and protein levels using Philosopher (v5.1.3-RC9). Identified peptides where quantified with DIA-NN (v1.8.2 beta 8) (Demichev et al., 2020) as implemented in the FragPipe package.

Peptide tables were filtered to retain only peptides quantified in at least 50% of replicates per condition for the N-termini enrichment experiment, and in at least 75% of replicates for the PICS experiment. Peptides exclusively observed in at least three replicates of one condition were likewise retained in both datasets. N-terminal peptides with a start position of ≤ 7 were excluded. Peptides sharing the same start position and N-terminal modification status but differing solely in methionine oxidation state or charge were collapsed into a single entry by summing their intensities. Differential abundance of N-terminal peptides between WT and *sbt12a-2* mutant samples was analyzed using the LIMMA package in R (Ritchie et al., 2015). Linear modeling with empirical Bayes moderation was applied to compute moderated *t*-statistics, p-values, and q-values, with log_2_ fold changes derived from replicate means. Changes in N-terminal peptide abundance supported by a LIMMA-moderated q-value < 0.05 were considered significant.

### Quenched peptide assays

The quenched peptides (QPs) derived from the candidate substrate protein were commercially synthesized with a DABCYL modification at the N-terminus and an EDANS modification at the C-terminus (GenScript) at a purity of 95%. The assay was performed as previously reported (Buscaill et al., 2024, Chen et al., 2024). Briefly, the synthesized peptides were resuspended in DMSO to prepare a 1 mM stock solution, which was further diluted in water to 200 µM. Immunoprecipitated SBT12a or SBT12a^S551A^ were mixed with QPs to a final concentration of 10 µM in a total volume of 100 µL. Fluorescence was recorded at 25°C for 2 hours using microplate reader (POLARstar Omega, BMG LABTECH, Ortenberg, Germany) with excitation and emission wavelengths set to 335 nm and 493 nm, respectively.

### Quantification and statistical analysis

All statistical analyses and generation of graphs were performed using GraphPad Prism software (version 9.5.1; GraphPad Software Inc.). The normality of the data was evaluated to determine the appropriate statistical method to be used. Brown–Forsythe and Welch ANOVA followed by Dunnett’s T3 multiple comparisons test was applied as parametric analyses. For non-parametric data, Kruskal–Wallis followed by Dunn’s multiple comparisons or Mann–Whitney tests were conducted. Sample size and statistical significance level have been indicated in the respective figure legends.

### Data availability

The mass spectrometry proteomics data have been deposited to the ProteomeXchange Consortium (Deutsch et al., 2023) via the PRIDE partner repository (Perez-Riverol et al., 2025) with the dataset identifiers PXD076390 for the HUNTER N-terminome data (Reviewer access token: hDUfTNzPSYEY) and PXD076264 for PICS data (Reviewer access token: xrYmY77Zs3Z2).

## Acknowledgements

We would like to thank Rosula Hinnenberg, Eija Schulze and Carmen Schubert for their excellent technical support, and the entire team for constructive discussions on the project and critical reviewing of the manuscript. We also acknowledge the staff of the Life Imaging Center (LIC) at the Hilde Mangold House (HMH), Albert-Ludwigs-University of Freiburg, for their support with confocal microscopy. Furthermore, we are grateful to Prof. Andreas Hiltbrunner for providing pPPO70v1HA plasmid. We thank Prof. Pascal Ratet for providing the Jemalong *dnf2* seeds. This work was supported by Enabling Nutrient Symbioses in Agriculture (ENSA) project, which was funded by Bill & Melinda Gates Agricultural Innovations (G1164932/57461) (P.M.D., T.O.), the German Research Foundation (DFG) under Germany’s Excellence strategy (CIBSS-EXC-2189 - project ID 390939984 to S.H., C.K., S.V.A., P.F.H., T.O.; SFB 1381 - project ID 403222702 to C.K., S.V.A., P.F.H., T.O.; SFB 1403 - project ID 414786233 to P.F.H.) and the China Scholarship Council (CSC) grant 202006300033 (G.Z.).

## Author contributions

Conceptualization, G.Z. and T.O.; Investigation, G.Z., F.S., M.M., M.R.F., F.V.B., C.H.R., B.L., W.Y., C.V.D.D., N.N., H.M.; Provided help in data analysis: A.M., S.H., N.N.; Supervision of individual project parts: C.K., C.S., P.M.D., P.F.H., T.O.; Writing-original draft, G.Z. and T.O.; Writing-review & editing, G.Z., F.S., M.M., M.R.F., F.V.B., W.Y., C.H.R., B.L., C.V.D.D., H.M., A.M., S.H., S.V.A., C.K., C.S., P.M.D., P.H., and T.O.; Funding acquisition, G.Z., P.M.D., P.F.H., and T.O.; Supervision, T.O.

## Declaration of interests

The authors declare that a patent application has been submitted (WO/2024/009146).

---

**Extended Data Fig. 1.**
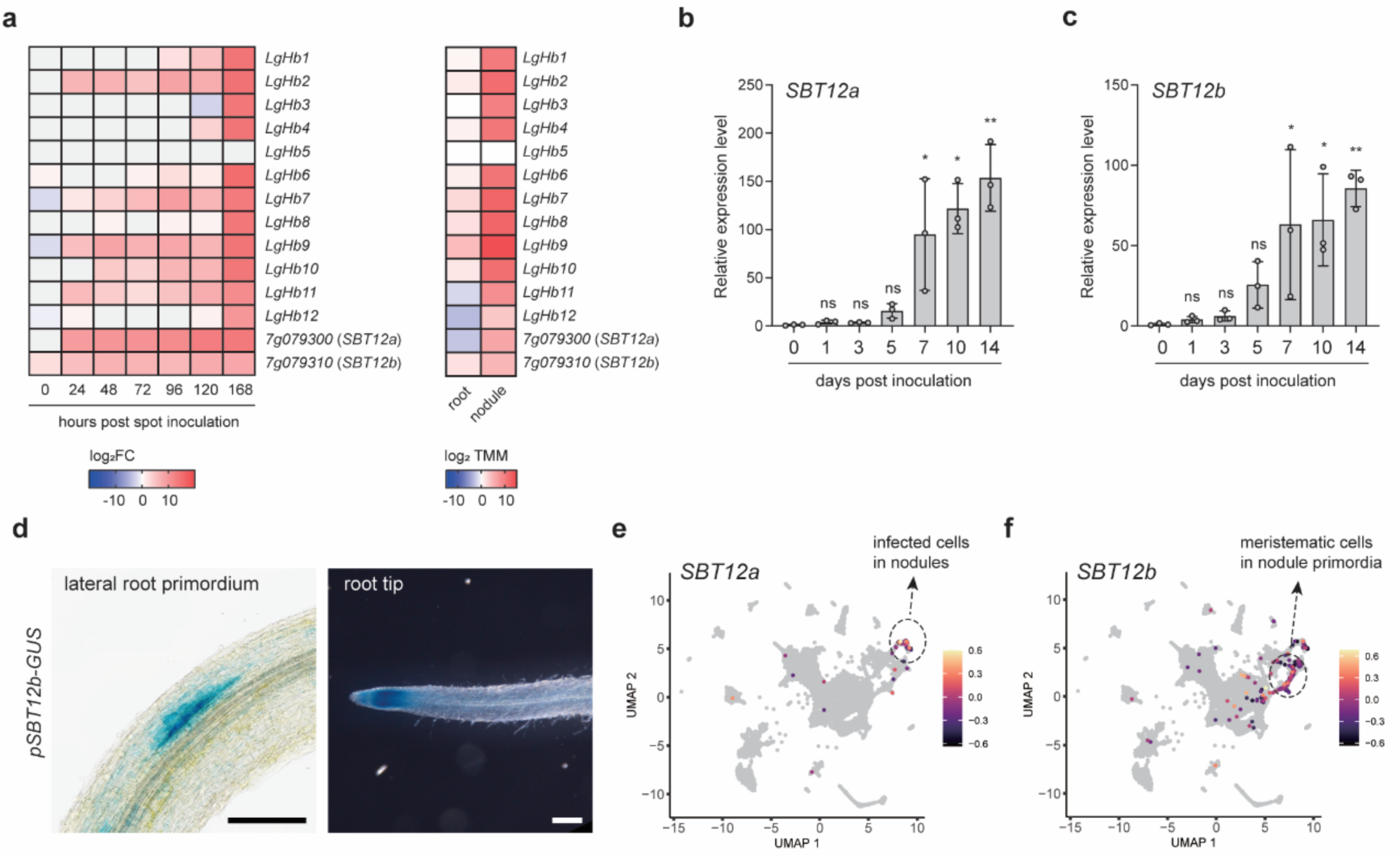
Expression analyses of the two subtilase genes, *SBT12a* and *SBT12b*, in *Medicago truncatula* during root nodule symbiosis. **a**, Left, expression heatmap showing the coordinated induction of 12 representative leghemoglobins (LgHbs) (Jiang et al., 2021) and two subtilase (SBT) genes, *Medtr7g079300* (*SBT12a*) and *Medtr7g079310* (*SBT12b*), during the nodulation process in Medicago (Schiessl et al., 2019). Right, expression comparison of *LgHb* and subtilase genes in nitrogen-starved WT roots and 10-day-old nodule tissues (Roux et al., 2014). **b,c**, qRT-PCR analysis of *SBT12a* (b) and *SBT12b* (c) expression in WT A17 roots inoculated with *S. meliloti 2011*. Root samples were harvested at indicated timepoints. Expression levels were normalized to ubiquitin. Values are the mean of three biological replicates and error bars represent standard deviations. Statistical differences were tested using Kruskal-Wallis test followed by Dunn’s multiple comparisons test. * p<0.05. ** p<0.01. ns indicates no significance (p>0.05). **d**, Activation of the *SBT12b* promoter in a lateral root primordium and a root tip of inoculated roots (5 dpi), visualized by a promoter-GUS assay. Scale bars, 200 μm. **e,f**, UMAP projection plots showing expression profiles of *SBT12a* (e) and *SBT12b* (f) across different cell types over the nodulation process (from 0 to 96 hpi). Cells that express *SBT12a* mainly belong to cluster 15, which contains infected cells that eventually become the nitrogen-fixing cells within the nodule. *SBT12b* expression was largely restricted to cells from cluster 9 that is composed of meristematic cells (Pereira et al., 2024).

**Extended Data Fig. 2.**
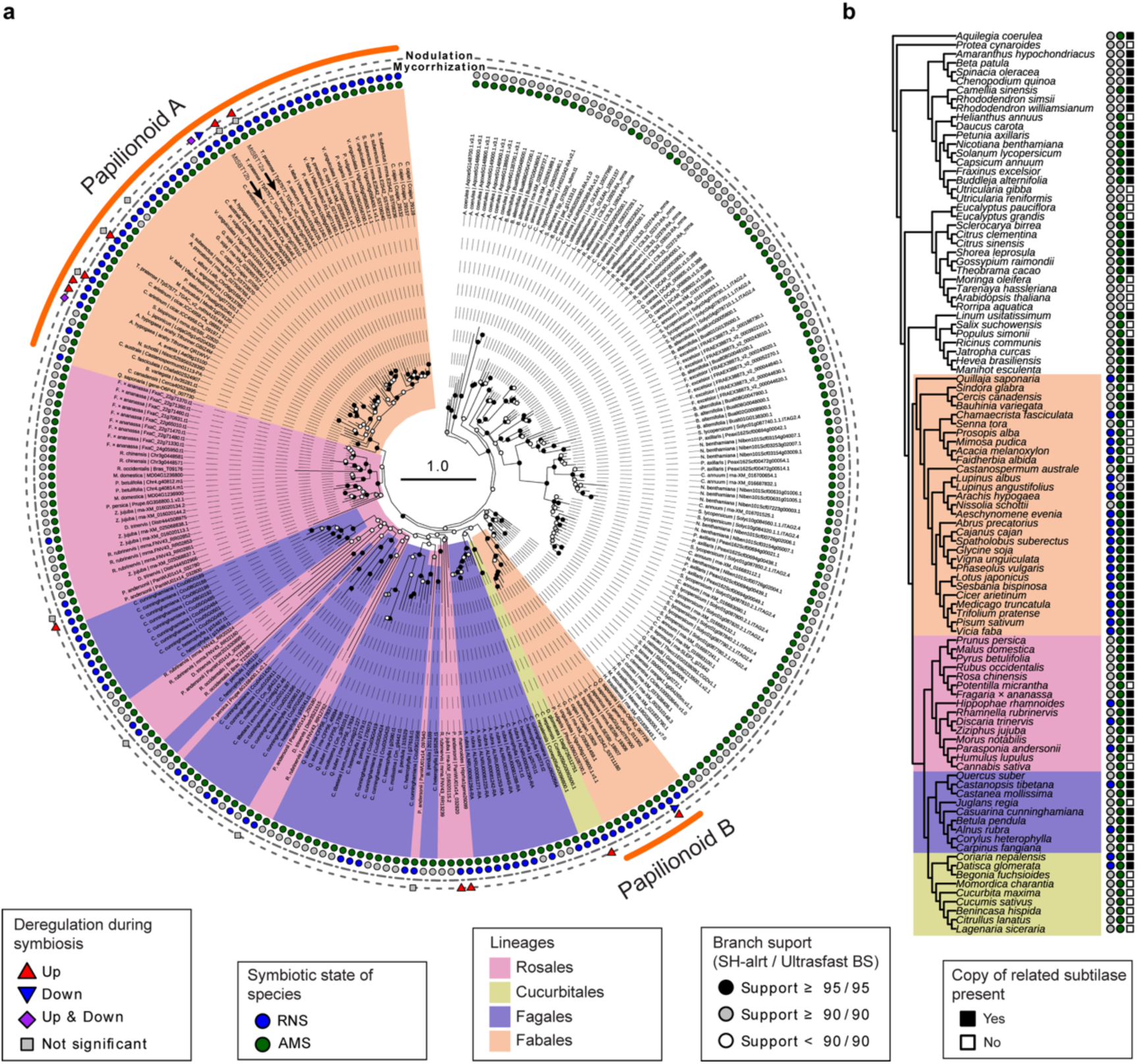
Phylogenetic analysis of subtilases related to SBT12a and SBT12b and their presence/absence in a variety of eudicots. **a**, Phylogenetic tree of subtilases reconstructed with maximum likelihood using the model “Q.plant+C60+F+I+R8”. Branch support is shown based on ultrafast bootstrap (% UFBS) and percentage of replicates of the SH-like approximate likelihood ratio test. **b**, Cladogram of eudicot species included in this analysis, with the presence or absence of the subtilases from (a). Symbiotic state of the species is shown for root nodule symbiosis (RNS; blue) and arbuscular mycorrhizal symbiosis (AMS; green). Genes known to be up- and/or downregulated during RNS or AMS based on the analysis from (a) are marked with a symbol on the outer circle of the tree. The two black arrows indicate the two genes from Medicago investigated in this study. The scale bar represents the estimated number of substitutions per site.

**Extended Data Fig. 3.**
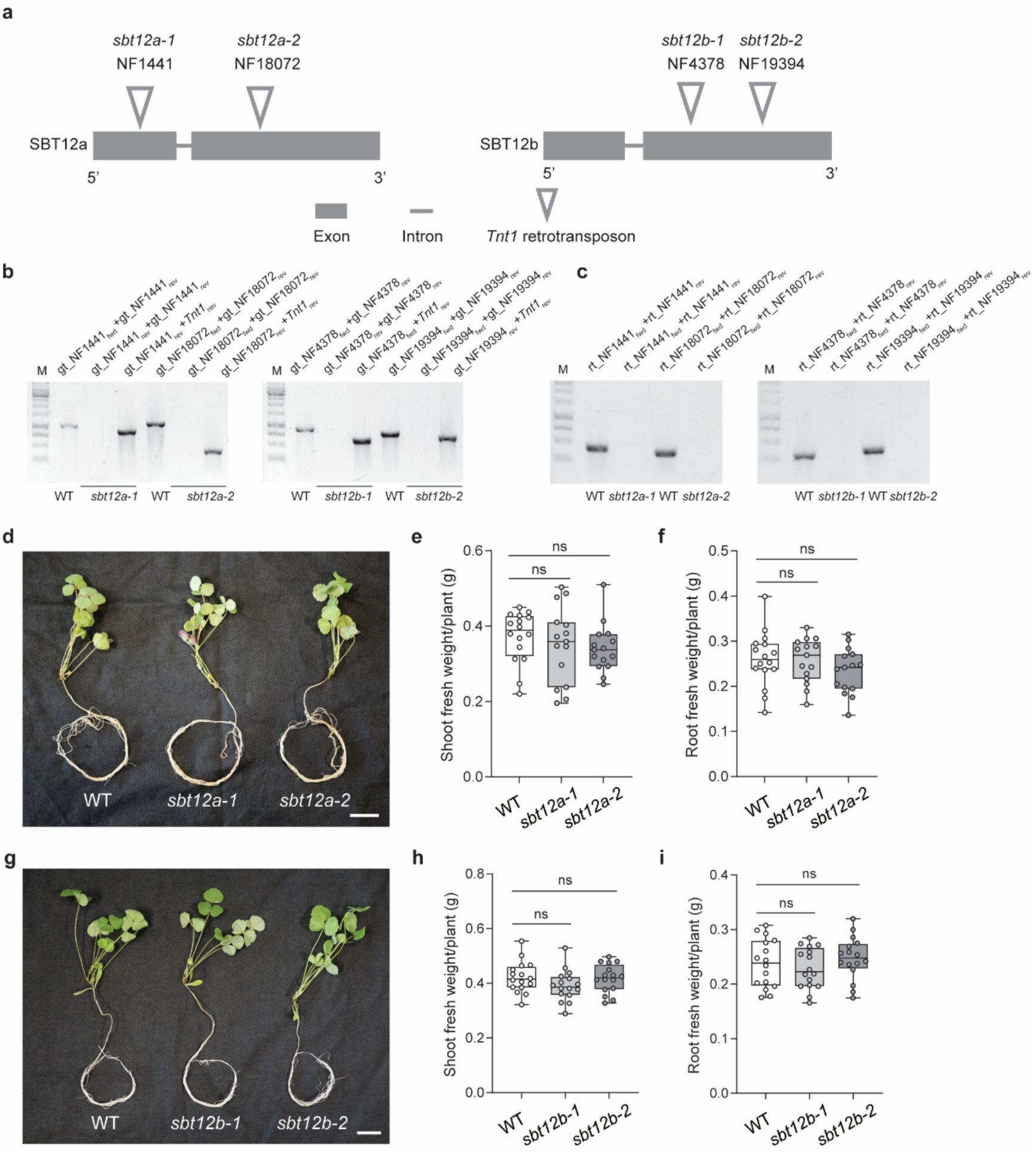
Characterization of *sbt12a-1*, *sbt12a-2*, *sbt12b-1* and *sbt12b-2* mutants in *Medicago truncatula*. **a**, Gene structure and positions of *Tnt1* retrotransposon insertions for *SBT12a* and *SBT12b*. **b**, Identification of *Tnt1* retrotransposon insertions in *sbt12a-1*, *sbt12a-2*, *sbt12b-1* and *sbt12b-2* mutant lines by PCR using genomic DNA as a template. For each line, gene specific primers flanking the putative insertion site were designed and named according to the corresponding insertion identifier. The primer binding the *Tnt1* retrotransposon was included to confirm the presence of the insertion fragment. **c**, RT-PCR analysis of *SBT12a* and *SBT12b* in *sbt12a* and *sbt12b* mutants, respectively. Primer pairs used in each lane are indicated above the agarose gels. M, marker. **d**, 28 days old *sbt12a* mutant seedlings grown in nitrogen-supplied conditions did not exhibit growth phenotypes compared to WT plants. Scale bar, 2 cm. **e**,**f**, Quantification of shoot (**e**) and root (**f**) fresh weight of *sbt12a* mutant and WT plants from (**d**). n=16, 15 and 15 for WT, *sbt12a-1* and *sbt12a-2*, respectively. Statistically significant differences were analyzed using Brown-Forsythe and Welch ANOVA test followed by Dunnett’s multiple T3 comparisons. ns indicates no significance (p>0.05). **g**, 28 days old *sbt12b* mutant plants grown in nitrogen-supplied conditions did not exhibit growth phenotypes compared to WT plants. Scale bar, 2 cm. **h**,**i**, Quantification of shoot (**h**) and root (**i**) fresh weight of *sbt12b* mutant and WT plants in (**g**). n=16 for all genotypes. Statistically significant differences were analyzed using Brown-Forsythe and Welch ANOVA tests followed by Dunnett’s multiple T3 comparisons. ns indicates no significance (p>0.05).

**Extended Data Fig. 4.**
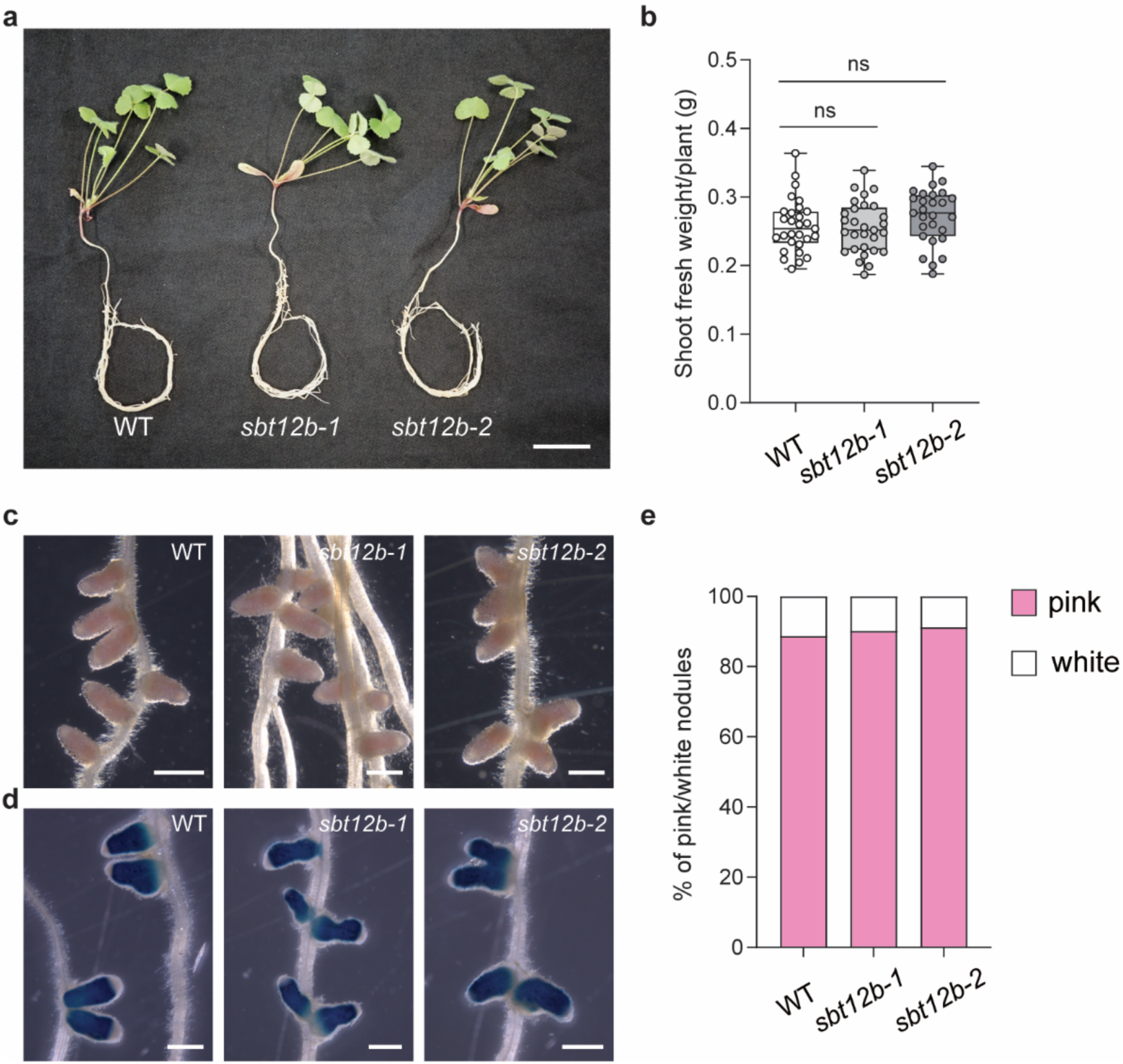
Phenotypic characterization of *sbt12b* mutants under symbiotic conditions. **a**, *sbt12b* mutant plants displayed no obvious difference compared to WT at 21 dpi with *S. meliloti 2011*. Scale bar, 2 cm. **b**, Quantification of shoot fresh weight of *sbt12b* mutant and WT plants at 21 dpi. n=29 for both WT and *sbt12b-1*, and 27 for *sbt12b-2*. Statistically significant differences were analyzed using Brown-Forsythe and Welch ANOVA test followed by Dunnett’s multiple T3 comparisons. ns represents no significance (p>0.05). **c**,**d**, Representative nodules formed on WT and *sbt12b* mutant roots at 21 dpi with *S. meliloti 2011 nifH::GUS*. No obvious differences in nodules developing on both *sbt12b* mutants were found in comparison to WT, as shown by the pink color of the nodules indicating that they are completely functional or overall activation of the GUS reporter indicating active nitrogen fixation. Scale bars, 1 mm. **e,** Ratio of pink and white nodules at 21 dpi. n=29 for both WT and *sbt12b-1*, and 27 for *sbt12b-2*. Nodulation phenotyping was independently repeated three times.

**Extended Data Fig. 5.**
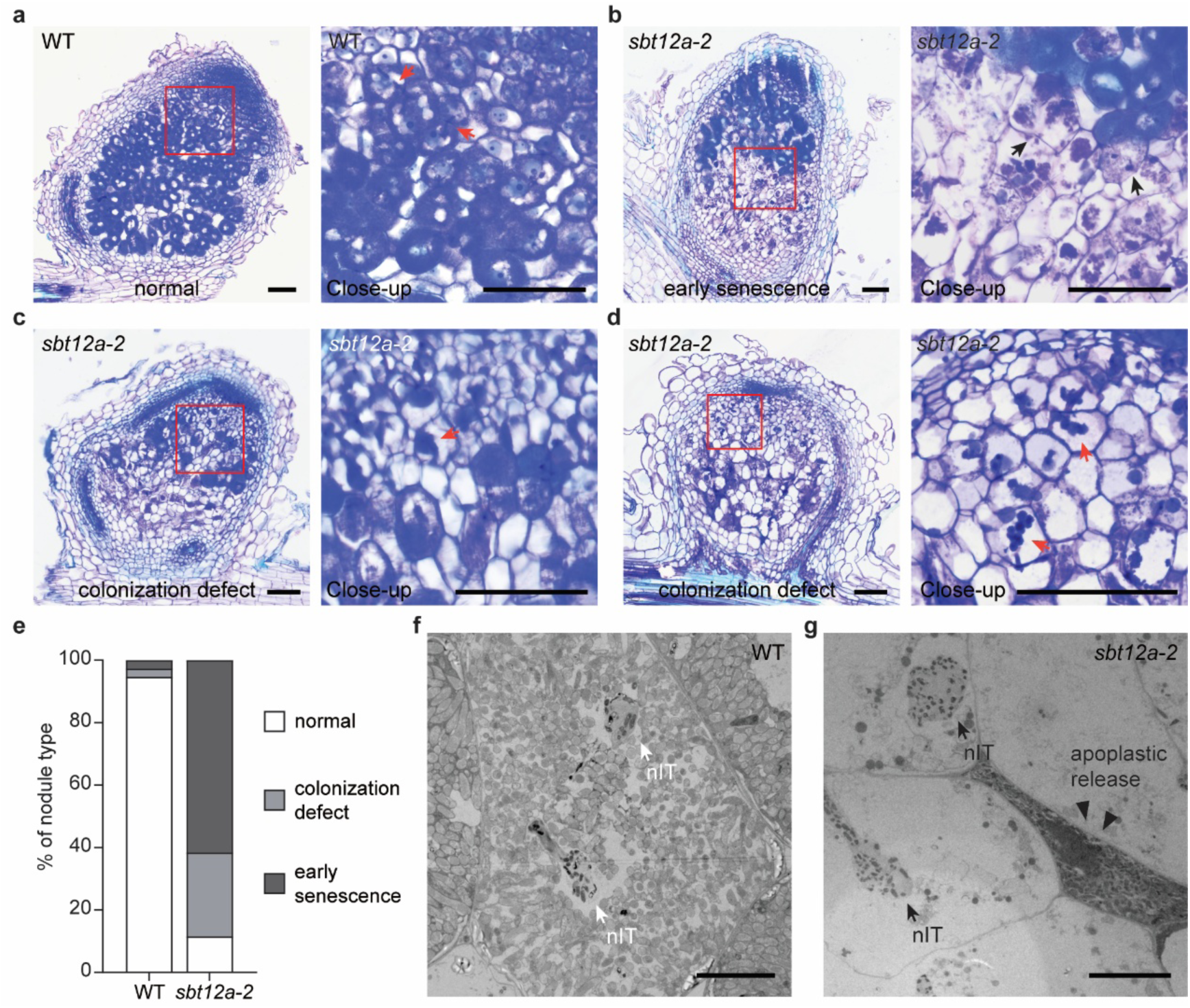
*SBT12a* is involved in nodule colonization. **a**-**d,** Longitudinal sections of WT (**a**) and *sbt12a-2* (**b**-**d**) nodules collected 14 dpi with *S. meliloti 2011*. Close-up views of the boxed regions are shown next to each panel. Note that in WT nodules, rhizobia were normally released from nodular infection threads (nITs, red arrows) and colonized host cells. In contrast, while the majority of *sbt12a-2* nodules show typical premature senescence phenotype (black arrows), about quarter of mutant nodules also exhibit rhizobial release defect, exemplified by swollen nITs. Scale bars, 100 μm. **e**, Quantification of WT and *sbt12a-2* nodule types as exemplified in **a**-**d**. 38 WT and 56 mutant nodules were randomly collected for sections from two independent replicates. **f**, TEM analysis showing representative rhizobial release sites in an infected WT nodule cell. **g**, *sbt12a-2* nodule cells displaying colonization defect, as indicated by enlarged nITs within largely uncolonized cells and apoplastic bacterial release. Scale bars, 10 μm (**f**, **g**).

**Extended Data Fig. 6.**
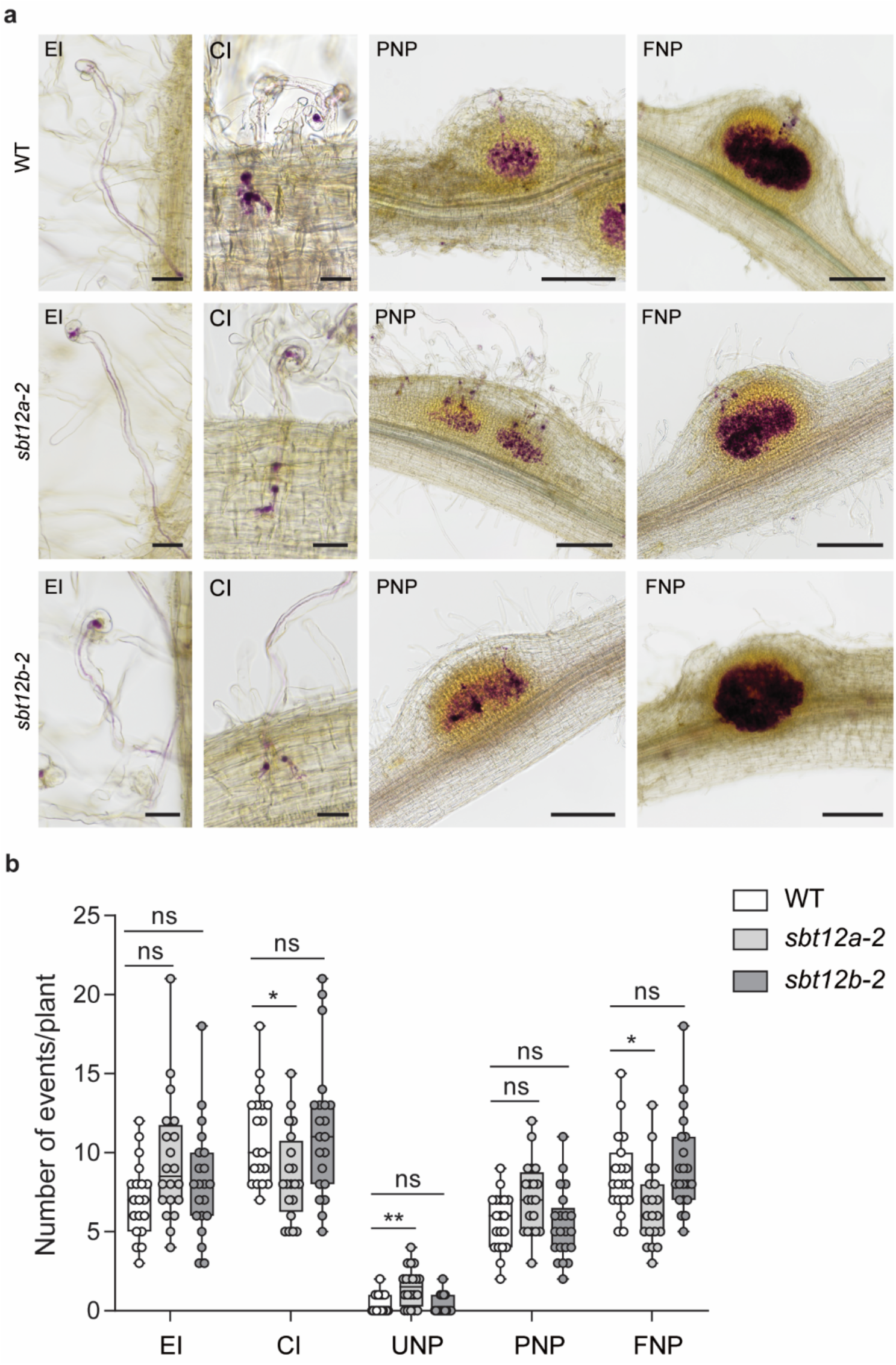
Phenotypic characterization of the *sbt12a* and *sbt12b* mutants during rhizobial infection. **a**, Representative images showing different stages of early nodule development in WT, *sbt12a-2* and *sbt12b-2* mutant. Roots were inoculated with *S. meliloti 2011* expressing *LacZ* (magenta stain) and harvested at 7 dpi for analysis. Infection events were categorized as followed: EI, epidermal infection; CI, cortical infection; PNP, partially infected nodule primordia. FNP, fully infected nodule primordia. UNP, uninfected nodule primordia. Scale bars, 30 μm (images classified as “EI” and “CI”) and 200 μm (images classified as “PNP” and “FNP”). **b**, Quantification of infection events from **a**. n=19, 20 and 21 for WT, *sbt12a-2* and *sbt12b-2*, respectively. Statistical significance was determined within each category using Kruskal-Wallis test followed by Dunn’s multiple comparisons test. * p<0.05. ** p<0.01. ns indicates no significance. The phenotyping experiment was repeated three times with similar results.

**Extended Data Fig. 7.**
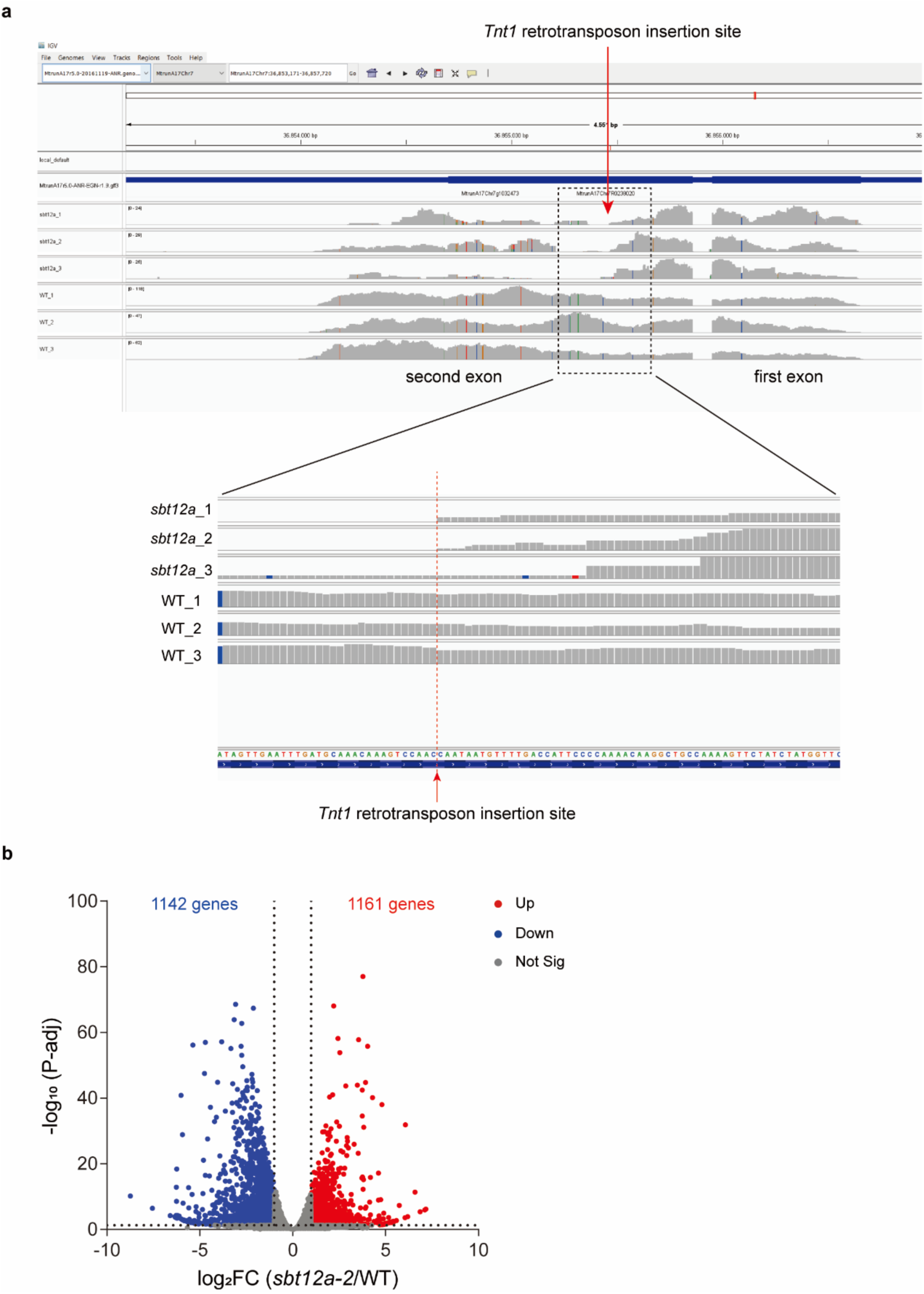
Transcriptional reprogramming by the *sbt12a* mutation. **a**, Distribution of reads mapping to the *SBT12a* gene from RNA-seq analysis. Three biological replicates for both genotypes are shown. Note that for all three *sbt12a-2* nodule samples, the read distribution after *Tnt1* retrotransposon insertion site (indicated by the red arrow) is completely disrupted. **b,** Volcano plot depicting the distribution of differentially expressed genes (DEGs) comparing *sbt12a-2* and WT nodules. Red and blue circles represent genes that are significantly upregulated (Up) and downregulated (Down) in *sbt12a-2* compared to WT R108, respectively (log2 fold change >1 and adjusted P-value<0.05). Grey circles represent genes that do not show significant changes (Not Sig) in expression levels.

**Extended Data Fig. 8.**
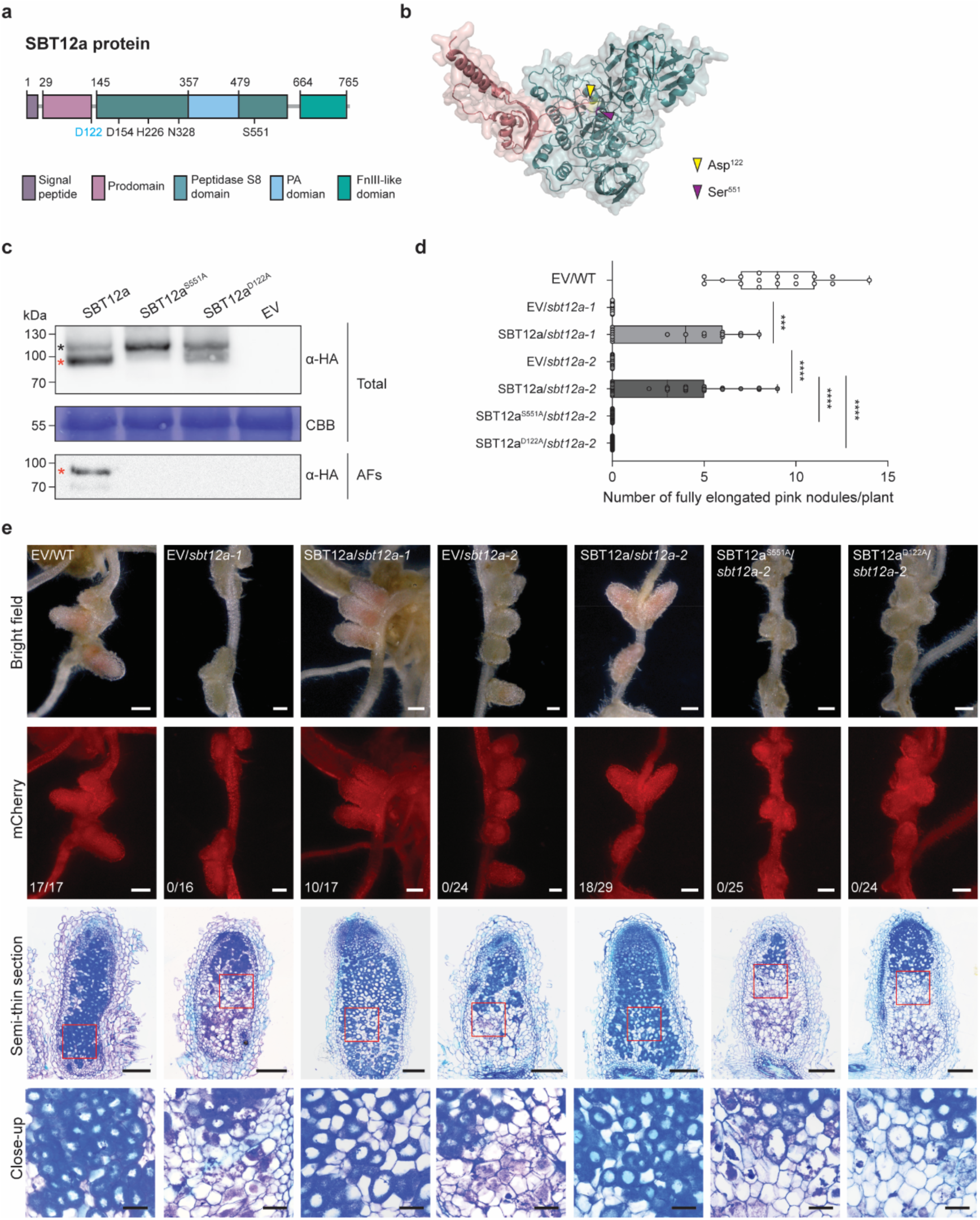
Characterization of SBT12a as a canonical subtilase. **a**, Schematic primary structure of the SBT12a protein. Different domains were predicted by InterPro (https://www.ebi.ac.uk/interpro/) and labeled with different colors. Black numbers indicate domain junctions and active amino acid residues. The Asp residue at the prodomain removal site was additionally labelled with blue. **b**, Modelled structure of the full-length SBT12a protein using AlphaFold3. Signal peptide and prodomain are colored in magenta and mature SBT12a in cyan. The proximity of the prodomain junction Asp-122 and the catalytic site Ser-551, indicated by yellow and purple arrowheads, respectively, suggests the potential self-processing during zymogen maturation. **c**, Immunoblot demonstrating the requirement of Ser-551 and Asp-122 for SBT12a self-processing and secretion. WT SBT12a and its mutant variants, S551A and D122A, were fused to the triple hemagglutinin (HA) tag and were transiently expressed in *N. benthamiana* leaves. An empty vector (EV) control was included. Leaf samples were harvested 4 days post agroinfiltration and subsequently subjected to total protein (Total) extraction or apoplastic fluids (AFs) isolation. Coomassie brilliant blue (CBB)-stained Rubisco proteins were used as a loading control. Black asterisk indicates SBT12a-3xHA precursor, and red asterisks indicate the mature form of SBT12a-3xHA after self-processing. **d**,**e**, Quantification of the number of pink nodules (**d**) and representative nodule phenotypes of transgenic roots transformed with empty vector (EV) or the indicated constructs in WT or *sbt12a* mutant backgrounds (**e**). Composite plants were inoculated with *S. meliloti 2011* and nodules were analyzed at 28 dpi. From top to bottom: bright-field images of representative nodules; images of mCherry fluorescence (red) fused to a nuclear localization signal (NLS), used as a selection marker for transformation; semi-thin nodule sections stained with toluidine blue; Close-up views of the boxed regions. Fully elongated pink nodules not displaying signs of premature senescence were considered as complemented nodules. The number of composite plants having complemented nodules relative to the total number of composite plants is indicated for each condition. Complementation experiments were independently repeated twice with similar results. Scale bars, 500 μm (bright field, mCherry channel); 200 μm (semi-thin sections) and 50 μm (close-up images). Statistical significance was determined using Mann-Whitney test. *** p<0.001. **** p<0.0001.

**Extended Data Fig. 9.**
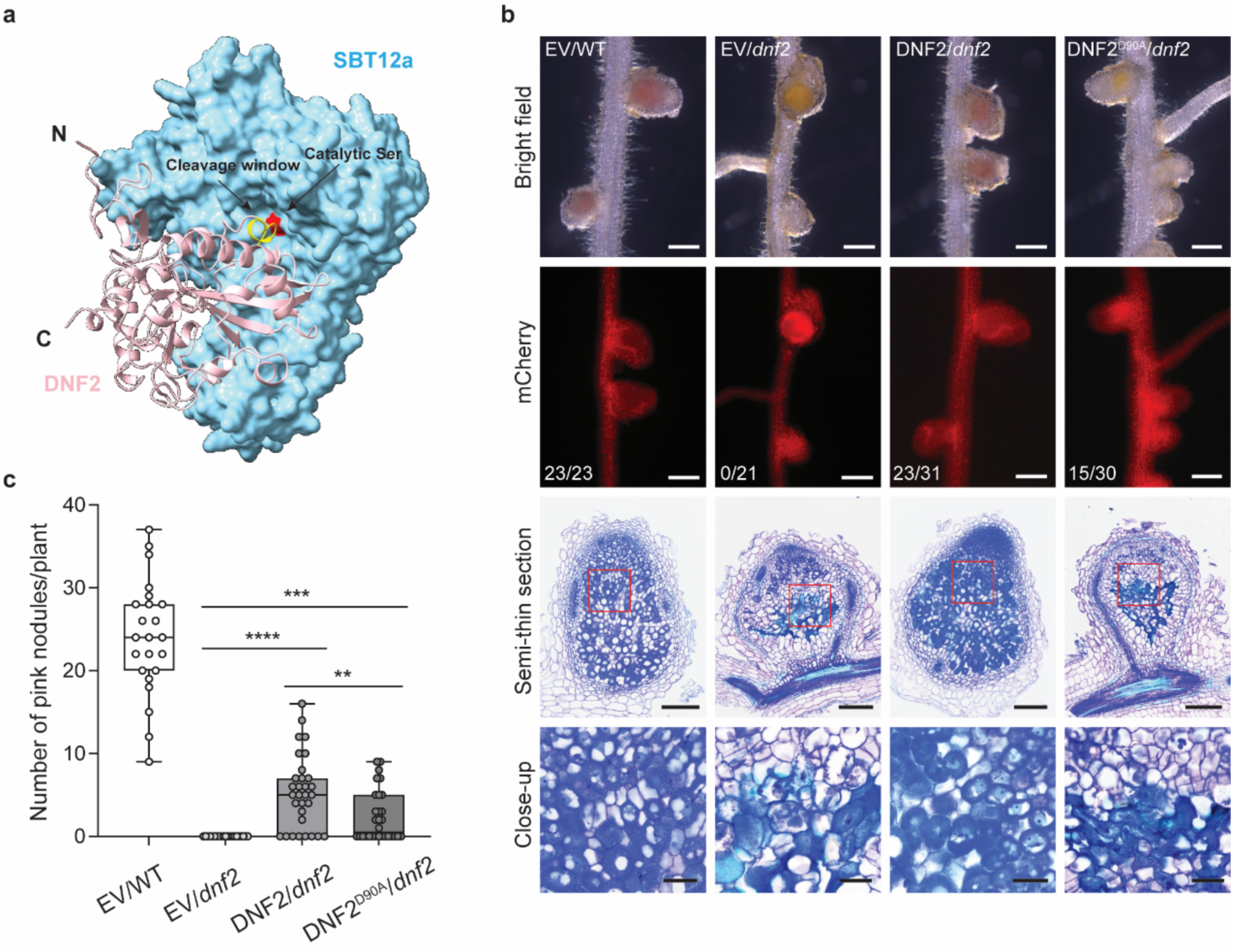
The D90 residue of DNF2 is important for its symbiotic function. **a**, Structural modelling of the SBT12a–DNF2 complex predicted by AlphaFold3 (ipTM+pTM=0.93), with SBT12a shown as a cyan surface and DNF2 as a pink cartoon. The N- and C-termini of DNF2 are indicated. The putative cleavage window within DNF2 (yellow) lies adjacent to the catalytic Ser of SBT12a (red). **b**, Genetic complementation of the *dnf2* mutant. Transgenic roots of WT or *dnf2* mutant transformed with the indicated constructs were inoculated with *S. meliloti 2011* and analyzed at 21 dpi. mCherry (red) fused to a NLS was used as a transgenic maker. Pink nodules not showing brown/yellow pigmentation (defense response) or signs of early senescence were considered as complemented nodules. The number of composite plants having complemented nodules relative to the total number of composite plants is indicated for each condition. Complementation experiments were independently repeated twice with similar results. Scale bars, 500 μm (bright field, mCherry channel); 200 μm (semi-thin sections) and 50 μm (close-up images). **c**, Quantification of the number of pink nodules per plant shown in **b**. Statistical significance was determined using the Mann–Whitney test. ** p<0.01. *** p<0.001. **** p<0.0001.

**Extended Data Fig. 10.**
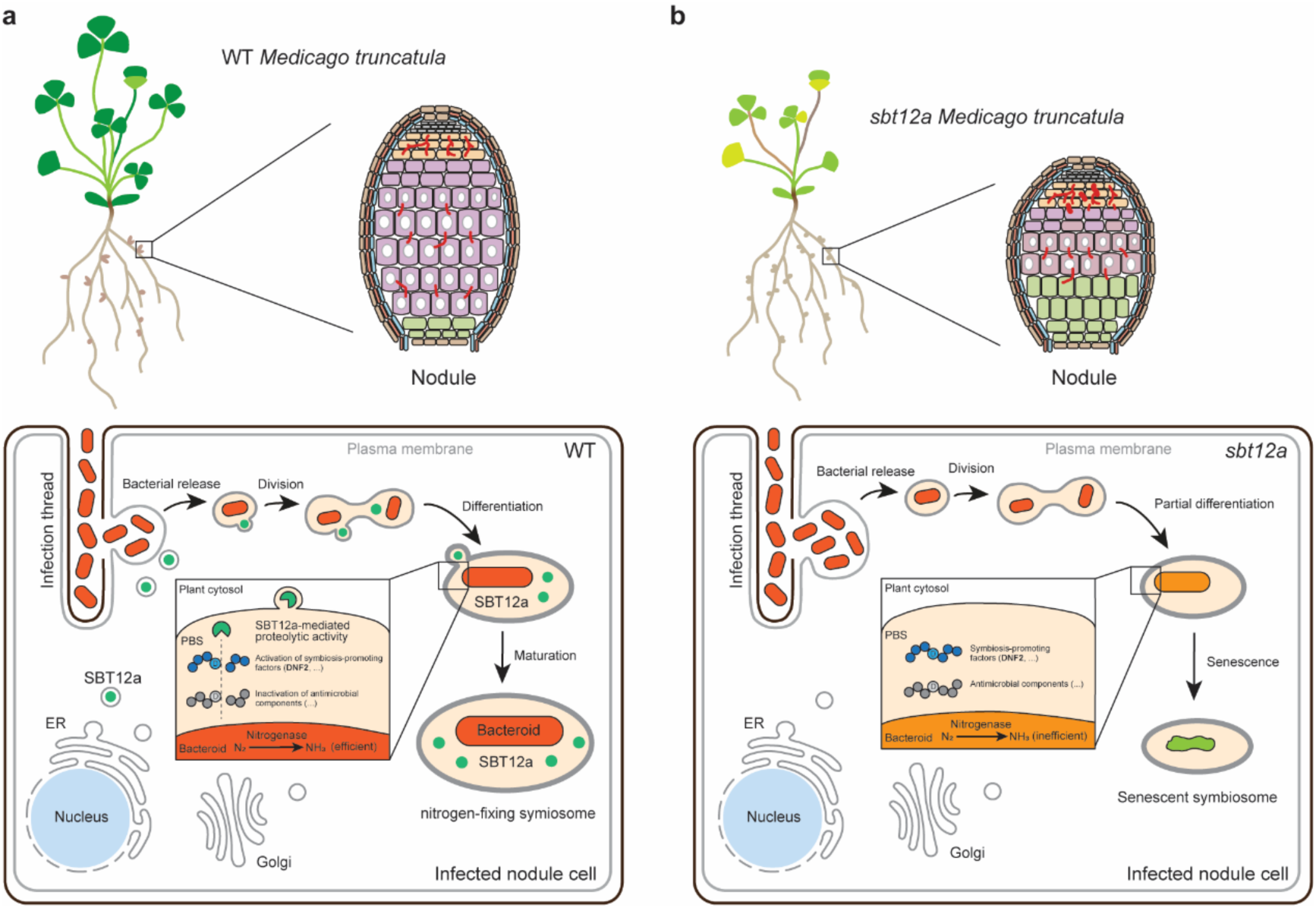
Proposed working model for SBT12a function. **a**, In WT nodules, rhizobia are released from infection threads into nodule cells, and subsequently enclosed within an organelle-like structure called the symbiosome. SBT12a is targeted to the bacterial release sites, and subsequently accumulates in newly formed symbiosomes. Within these symbiosomes, rhizobia proliferate and differentiate into a special form known as bacteroids. This developmental transition is accompanied by the sustained SBT12a-mediated proteolysis in the peribacteroid space. Acting there as a phytaspase with Asp-specific cleavage specificity, SBT12a drives symbiosome stabilization, which is indispensable for sufficient nitrogen fixation occurring in bacteroids. The activity of SBT12a likely involves the modulation of symbiosis-promoting regulators, such as DNF2, or the degradation of antimicrobial components. **b**, In the *sbt12a* mutant, the absence of SBT12a-mediated proteolytic regulation within the peribacteroid space leads to severe defects in bacteroid differentiation. Consequently, impaired symbiosome maintenance fails to support efficient nitrogen fixation, leading to premature nodule senescence and nitrogen starvation in seedlings.

## Notes

### Summary of Updates

In the revision, we have mainly added results for the characterization of the SBT12a target DNF2. These results can be found in the new Fig. 6 and Extended Data Fig 9. In addition, we have added additional information requested by the reviewers.

